# Disulfide stabilization reveals conserved dynamic features between SARS-CoV-1 and SARS-CoV-2 spikes

**DOI:** 10.1101/2022.10.14.512296

**Authors:** Xixi Zhang, Zimu Li, Yanjun Zhang, Yutong Liu, Jingjing Wang, Banghui Liu, Qiuluan Chen, Qian Wang, Lutang Fu, Peiyi Wang, Xiaolin Zhong, Liang Jin, Ling Chen, Jun He, Jincun Zhao, Xiaoli Xiong

**Author notes:** These authors contribute equally (Xixi Zhang, Zimu Li, Yanjun Zhang, Yutong Liu).

## Abstract

SARS-CoV-2 spike protein (S) is structurally dynamic and has been observed by cryo-EM to adopt a variety of prefusion conformations that can be categorized as locked, closed and open. The locked conformations feature tightly packed trimers with structural elements incompatible with RBD in “up” position. For SARS-CoV-2 S, it has been shown that the locked conformations are transient under neutral pH. Probably due to their transience, locked conformations remain largely uncharacterized for SARS-CoV-1 S. Intriguingly, locked conformations were the only conformations captured for S proteins of bat and pangolin origin SARS-related coronaviruses. In this study, we introduced x1, x2, and x3 disulfides into SARS-CoV-1 S. Some of these disulfides have been shown to preserve rare locked conformations when introduced to SARS-CoV-2 S. Introduction of these disulfides allowed us to image a variety of locked and other rare conformations for SARS-CoV-1 S by cryo-EM. We identified bound cofactors and structural features that are associated with SARS-CoV-1 S locked conformations. We compare newly determined structures to other available spike structures of Sarbecoviruses to identify conserved features and discuss their possible functions.

## Introduction

Betacoronavirus subgenus *Sarbecovirus* is composed of SARS-CoV-1, SARS-CoV-2 and other severe acute respiratory syndrome-related coronaviruses (SARSr-CoVs) (Coronaviridae Study Group of the International Committee on Taxonomy of Viruses 2020). Due to SARS-CoV-1’s known ability to infect humans and the SARS-CoV-2 pandemic, Sarbecoviruses are subjected to intense research. Among structural proteins of Sarbecoviruses, spike (S) protein is responsible for receptor binding and mediates membrane fusion between cell and virus to initiate infection (Li, 2016). Being the most exposed protein on the virus surface, it is the main target of immune system. Most available SARS-CoV-2 vaccines utilize S protein or its derivatives as the immunogen (Krammer, 2020) and a number of antibodies targeting the SARS-CoV-2 S protein have been approved for emergency use authorization (Corti *et al*, 2021). For these reasons, S protein is a major focus for coronavirus vaccine and therapeutics development.

Murine Hepatitis Virus (MHV) (Walls *et al*, 2016a), human coronavirus HKU1 (HCoV-HKU1) (Kirchdoerfer *et al*, 2016) and human coronavirus NL63 (HCoV-NL63) (Walls *et al*, 2016b) spikes were the first structurally characterized coronavirus (CoV) spikes. They all adopt single conformations with all their 3 receptor binding domains (RBDs) in “down” positions. Among Sarbecoviruses, structures of SARS-CoV-1 S protein were first determined and conformational dynamics was first observed for CoV spikes. RBDs in SARS-CoV-1 spikes were observed to be in “up” and “down” positions making the spikes exhibit open and closed conformations respectively (Gui *et al*, 2017; Yuan *et al*, 2017). In line with SARS-CoV-1 S, RBD “up” open and RBD “down” closed spikes were observed when SARS-CoV-2 S structures were initially determined (Walls *et al*, 2020; Wrapp *et al*, 2020). However, a third prefusion conformation, called “locked”, was subsequently identified in multiple studies using either full-length spikes or spike ectodomains (Bangaru *et al*, 2020; Cai *et al*, 2020; Toelzer *et al*, 2020; Wrobel *et al*, 2020; Xiong *et al*, 2020). In pH neutral phosphate buffered saline (PBS), this conformation appeared to be transient and only a small fraction of purified spikes adopted locked conformations (Xiong *et al*., 2020). Overall, locked conformations are more ordered, adopting a more tightly packed trimeric quaternary structure, possessing features including, bound lipid in RBD, rigidified Domain D region and ordered fusion peptide proximal region (FPPR, residues 833-855 in SARS-CoV-2 S).

Detailed structural analysis of SARS-CoV-2 S structures identified that interactions between Domain D and Domain C-D hinge region in locked conformations likely restrain RBD movement leading to more compact spike packing (Qu *et al*, 2022). A comparison to other CoV S structures revealed that HCoV-NL63 (Walls *et al*., 2016b), MHV (Walls *et al*., 2016a), Infectious Bronchitis Virus (IBV) (Shang *et al*, 2018) and Porcine Delta coronavirus (PDCoV) (Xiong *et al*, 2018) spikes are structurally more related to locked structures of SARS-CoV-2 spike, they all have ordered FPPR which is incompatible with RBD in “up” position (Xiong *et al*., 2020). More perplexingly, locked conformations were not observed for spikes on fixed SARS-CoV-2 virus particles (Ke et al, 2020; Turoňová et al, 2020; Yao et al, 2020).

We and others engineered disulfide bonds to trap SARS-CoV-2 spike in RBD “down” conformations (Henderson *et al*, 2020; McCallum *et al*, 2020; Xiong *et al*., 2020). In addition to increased spike stability, we found that x1 (Xiong *et al*., 2020) and x3 (Qu *et al*., 2022) disulfide bonds increase proportions of purified spikes adopting locked conformations. Structural studies of x3 disulfide stabilized locked spikes allowed us to further classify locked spikes into “locked-1” and “locked-2” conformations, which differ primarily in ways how Domain D region rigidifies. Protomers in a SARS-CoV-2 S trimer can adopt exclusively “locked-1” or “locked-2” conformations or a trimer can be formed by protomers of the two locked conformations in various combinations (Qu *et al*., 2022). Together with “closed” and “open” conformations, prefusion SARS-CoV-2 spike exhibits complex dynamics.

Despite considerable sequence homology to SARS-CoV-2 S, currently determined structures of SARS-CoV-1 spikes are either in closed or open conformation (Gui *et al*., 2017; Yuan *et al*., 2017). Locked conformations remain elusive for SARS-CoV-1 spike, therefore dynamics of the spike likely remains incompletely characterized. In this study, we engineered SARS-CoV-1 spikes bearing the x1, x2, and x3 disulfides. We purified engineered SARS-CoV-1 spikes and characterized their biochemical properties. By Cryo-EM, engineered SARS-CoV-1 spikes allowed us to determine a series of spike structures adopting previously undetected conformations. We compared these structures to available spike structures of other Sarbecoviruses to identify conserved features.

### Production of SARS-CoV-1 spikes with engineered disulfide bonds

x1, x2 and x3 (sites in SARS-CoV-2 S numbering: x1, S383C and D985C; x2, G413C and V987C; x3, D427C and V987C) disulfide bonds were designed for SARS-CoV-2 spike (Qu *et al*., 2022; Xiong *et al*., 2020). These disulfides facilitate formation of covalently linked spike trimers through the introduced disulfides between RBD and S2 of a neighboring spike protomer. x1 disulfide has been previously introduced in SARS-CoV-1 spike to show successful disulfide formation (McCallum *et al*., 2020). We introduced these three disulfide bonds in equivalent positions in the SARS-CoV-1 S sequence generating three spike variants: S/x1 (S370C and D967C), S/x2 (G400C and V969C) and S/x3 (D414C and V969C) (**Fig. 1a** and **Fig. S1**). These spike variants were expressed in Expi293 cells to be secreted into culture media. Compared to purified unmodified SARS-CoV-1 spike ectodomain, purified S/x1, S/x2 and S/x3 SARS-CoV-1 spikes successfully form covalently linked trimers in SDS-PAGE under non-reducing conditions and these trimers could be reduced to monomers under denaturing conditions (**Fig. 1b and Fig. S2**). x1, x2, and x3 disulfides in SARS-CoV-1 spike exhibit different sensitivity to reducing agent. Under native conditions, x2 disulfide is more resistant to reduction by DTT than x1 and x3 disulfides, likely reflecting different local chemical environments for different disulfides (**Fig. S2**). Resistance to DTT reduction was also observed for x2 disulfide in SARS-CoV-2 spike (Qu *et al*., 2022). Negative-staining EM of purified spike variants showed that S/x1, S/x2 and S/x3 spikes form trimers with morphologies indistinguishable to unmodified SARS-CoV-1 spike ectodomain (S/native) (**Fig. 1c**). These results confirmed that x1, x2 and x3 disulfides previously designed for SARS-CoV-2 spike are compatible with production of well-formed covalently linked SARS-CoV-1 spike trimers.

**Fig. 1.**
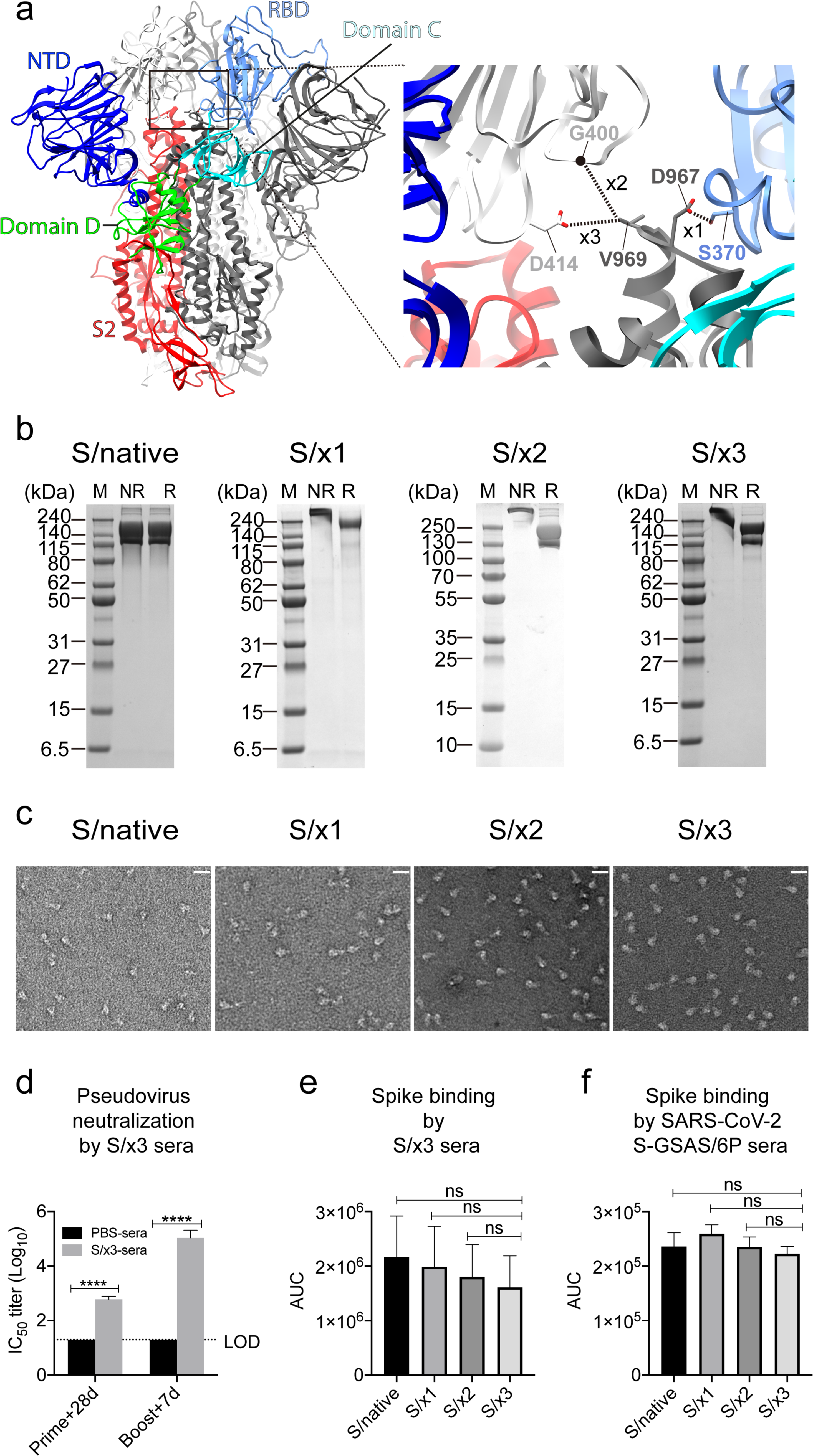
Design, purification and characterization of SARS-CoV-1 S/x1, S/x2 and S/x3 spikes. **a**, Structure of SARS-CoV-1 spike (PDB: 5X58 (Yuan *et al*., 2017)) in closed conformation (left panel). The box indicates the location of the zoomed-in view in the right panel, engineered x1 (S370C and D967C), x2 (G400C and V969C) and x3 (D414C and V969C) disulfides are indicated in the zoomed-in view. **b**, Coomassie-stained SDS-PAGE gels of purified SARS-CoV-1 S/native, S/x1, S/x2 and S/x3 spikes. NR or R indicates protein sample prepared in either non-reducing or reducing condition. M indicates protein marker lanes. **c**, Negative stain EM images of the purified spikes. The white lines indicate a length of 250 Å. **d**, Immunogenic and antigenic characteristics of SARS-CoV-1 spikes. Left panel, SARS-CoV-1 pseudovirus neutralizing titers of S/x3 primed and boosted mouse immune sera. Mean and SEM (error bar) are shown, dashed line represents limit of detection (LOD). \*\*\*\**P* < 0.0001, *P* values were analyzed by 2-way ANOVA test. Middle panel, binding of S/native, S/x1, S/x2 and S/x3 spikes by the S/x3 boosted mouse immune sera. Right panel, binding of S/native, S/x1, S/x2 and S/x3 spikes by the SARS-CoV-2 S-GSAS/6P boosted mouse immune sera. Binding by immune sera was measured by areas under the curves (AUC) derived from ELISA titration curves (**Fig. S3**). Mean and SEM (error bar) are shown. ns, not significant, *P* values were analyzed by one-way ANOVA test.

### Antigenic and immunogenic properties of engineered spikes

Purified S/x3 spike was used to immunize mice employing a prime-boost immunization strategy. S/x3 was found to exclusively adopt an RBD “down” locked-1 conformation by cryo-EM (**Fig. 2h**). After priming by 10 µg of S/x3 spike, mouse sera showed induction of neutralization titers against SARS-CoV-1 pseudovirus (**Fig. 1d**). Boost by another dose of 10 µg of S/x3 spike, increased neutralization titer further by approximately 100-fold (**Fig. 1d**). The complete immunization regimen induced a serum neutralization titer of approximately 5 log10 dilution units (**Fig. 1d**). We found mouse immune sera raised with engineered SARS-CoV-2 RBD “down” S-R/x2 (“-R” stands for furin site changed to a single “arginine” and “x2” stands for the introduced x2 disulfide) spike protein as the immunogen gave similar neutralization titers (Carnell *et al*, 2021). By ELISA (enzyme-linked immunosorbent assay), the S/x3 induced immune sera exhibit binding towards SARS-CoV-1 S/native, S/x1, S/x2, and S/x3 spikes. There are differences in reactivity of S/x3 immune sera towards different spikes, but the differences are not statistically significant (**Fig. 1e, Fig. S3a**). SARS-CoV-2 S-GSAS/6P immune sera cross-react with S/native, S/x1, S/x2, and S/x3 spikes, and as expected, it showed reduced reactivity compared to S/x3 immune sera (**Fig. 1f, Fig. S3b**).

**Fig. 2.**
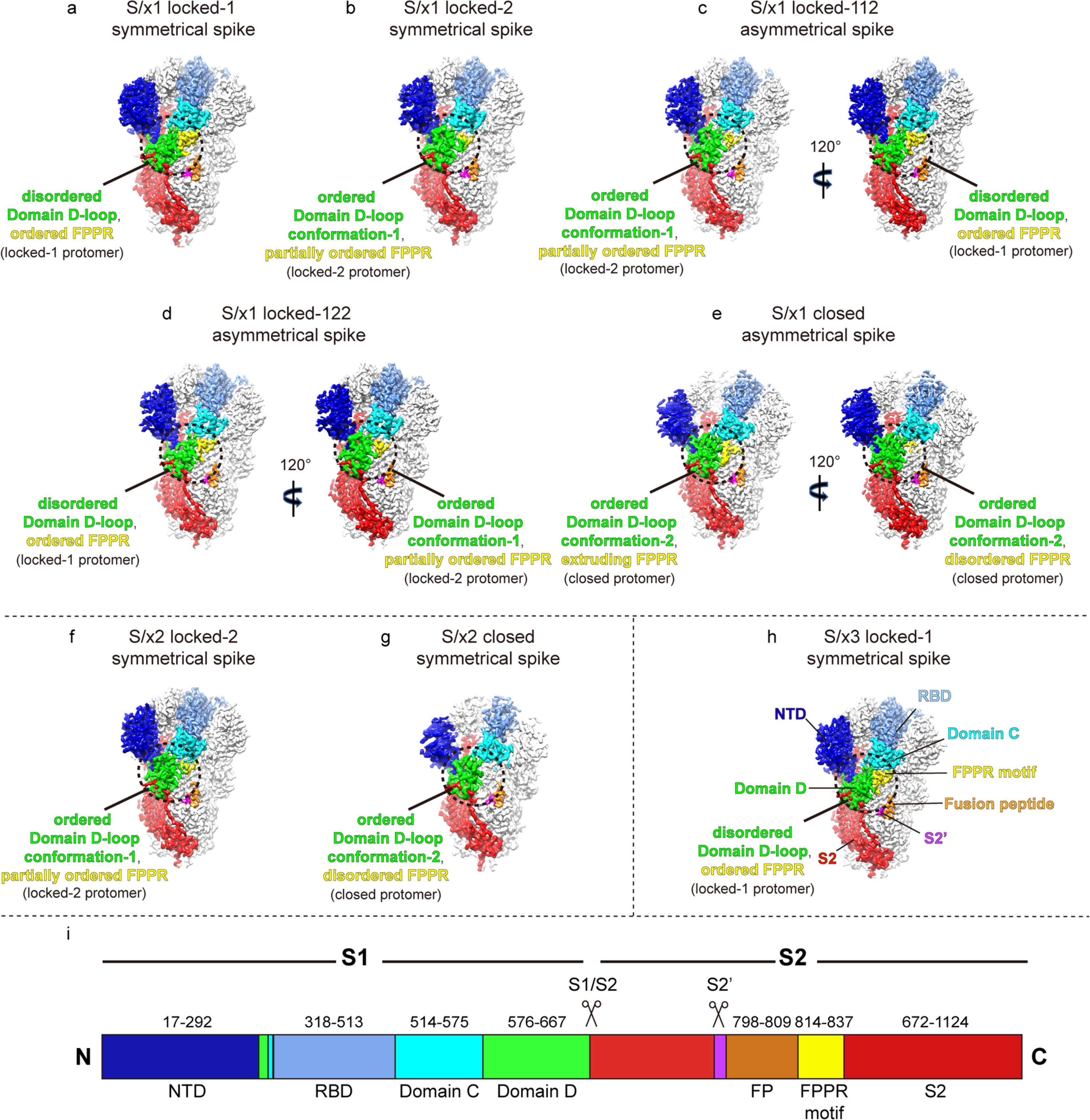
Cryo-EM structures of SARS-CoV-1 S/x1, S/x2 and S/x3 spikes reveal multiple spike conformations. **a-h**, Cryo-EM densities are shown for structures determined for SARS-CoV-1 S/x1, S/x2 and S/x3 spikes. Featured protomers within the spike trimer densities are colored. S1 structural domains - NTD, RBD, Domain C, and Domain D are colored blue, light blue, cyan, and green respectively. S2 is colored red with S2 structural elements - S2’, FP (fusion peptide) and FPPR (fusion peptide proximal region) highlighted magenta, orange and yellow respectively. **a-b**, Symmetrical locked-1 and locked-2 conformations identified for S/x1. S/x1 locked-1 conformation has a disordered Domain D-loop and ordered FPPR. Whereas Domain D-loop in S/x1 locked-2 conformation adopts an ordered conformation (Domain D-loop conformation-1) and FPPR is partially ordered. Domain D regions where major structural rearrangement occurs are highlighted within the dashed circles. **c-d**, Asymmetrical locked-112 and locked-211 conformations identified for S/x1 spike. Spike trimers in these conformations are assembled from different combinations of locked-1 and locked-2 protomers. Spike trimers are rotated by 120° to show protomers of different conformations. **e**, S/x1 spike in an asymmetrical closed conformation. The spike is rotated 120° to show structural differences in FPPR between protomers. **f-h**, Symmetrical conformations identified for S/x2 and S/x3 spikes. **i**, A schematic drawing of SARS-CoV-1 spike sequence showing its structural domains and elements.

### SARS-CoV-1 spikes with engineered disulfide bonds reveal additional conformations

We determined cryo-EM structures of SARS-CoV-1 S/x1, S/x2 and S/x3 spikes and we identified various previously undetected conformations for SARS-CoV-1 S through classification procedures (see **Fig. S4**). SARS-CoV-1 S/x1 spike exhibited complex dynamics, we observed spikes adopting both symmetrical and asymmetrical conformations. Symmetrical locked-1 (3 locked-1 protomers) and symmetrical locked-2 (3 locked-2 protomers) spike trimers differ at Domain D-loop (residues 602-627) and FPPR structures (**Fig. 2a, b** and **Fig. 3a, b**). In addition, we observed asymmetrical locked-112 (2 locked-1 protomers and 1 locked-2 protomer, **Fig. 2c**) and locked-122 (1 locked-1 protomer and 2 locked-2 protomers, **Fig. 2d**) spike trimers. Similar structural heterogeneity has been observed for the SARS-CoV-2 S-R/x3 spike in various locked conformations (Qu *et al*., 2022). Interestingly, SARS-CoV-1 S/x1 spike also adopts an asymmetrical closed conformation: FPPR in one protomer adopts an ordered extruding conformation while being disordered in the other two protomers (**Fig. 2e, Fig. 3d**). By cryo-EM, only symmetrical spike trimers were observed for SARS-CoV-1 S/x2 and S/x3 spikes. S/x2 spike adopts symmetrical locked-2 (**Fig. 2f** and **Fig. 3c**) and closed (**Fig. 2g** and **Fig. S5a**) conformations. S/x3 spike was only observed to adopt a symmetrical locked-1 conformation (**Fig. 2h, Fig. S5b**). Overall, locked-1, locked-2 are structurally more rigid than closed conformations showing tighter packing and less flexibility towards RBD end of spike (**Fig. S6 and Fig. S7**). Structures within each identified conformation categories, namely locked-1, locked-2 and closed, share almost identical structural features. Between different conformation categories, there are variations in structures of Domain D, FPPR and positioning of RBD. These variations appear to result different spike trimer packing (**Fig. S7**). Distinct structures from different conformation categories are summarized and described in details in the next sections.

**Fig. 3.**
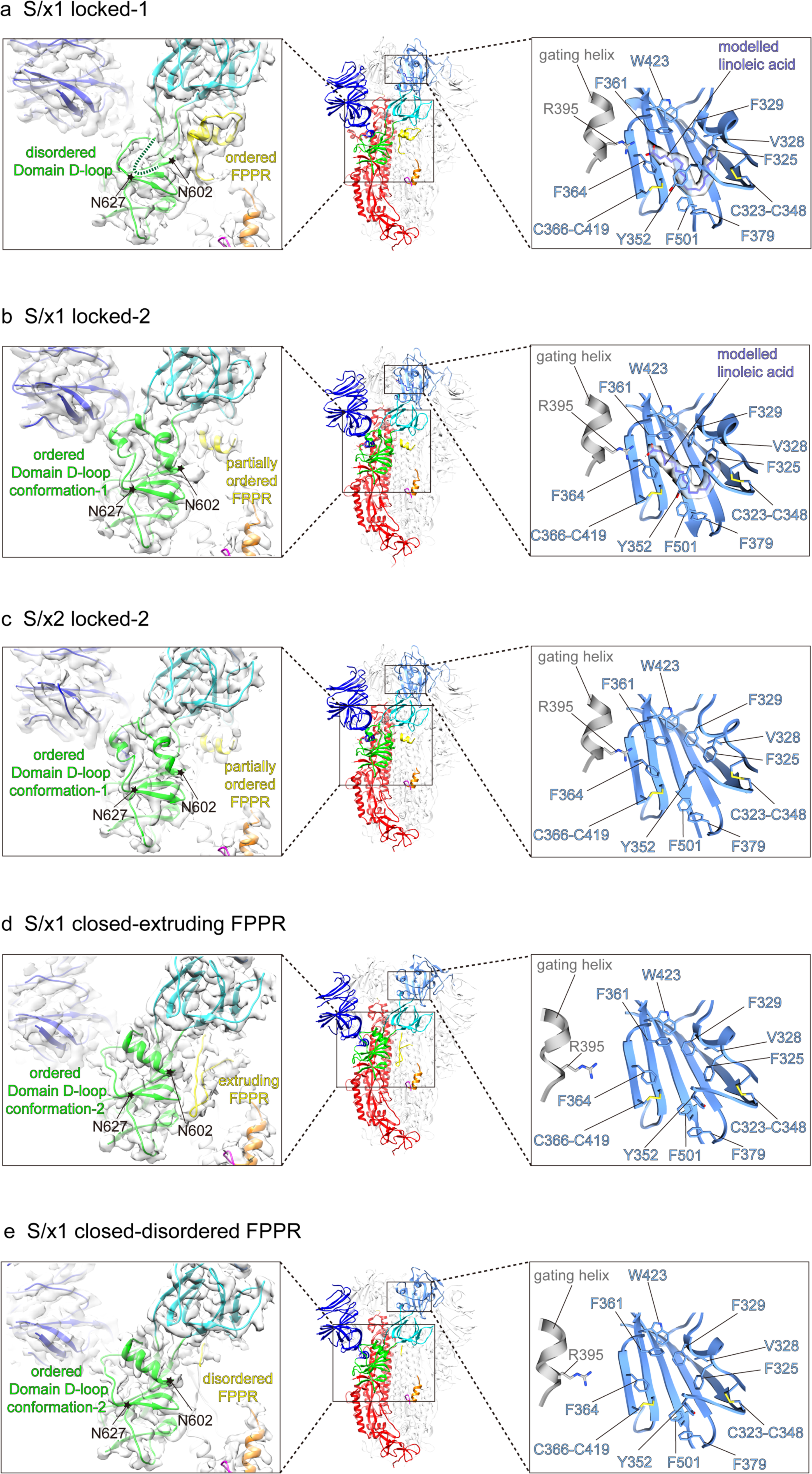
Structural features of spike protein protomers in different conformations. **a-e**, Representative protomers of different conformation categories and with distinctive features were summarized from structures shown in Fig. 2. Molecular models are shown in Middle panels. Left panels show cryo-EM densities of Domain D and surrounding regions with fitted molecular models. Flexible regions (Domain D-loop) between N602-N627 where structural rearrangements occur are highlighted and indicated by star symbols. Structural features in each structure are shown and indicated. Right panels show lipid binding pockets within SARS-CoV-1 RBDs of different conformations. Hydrophobic amino acid sidechains forming the lipid binding pocket are shown in stick representations. Gating helices from the neighboring RBDs are shown in grey with the head group interacting R395 in stick representations. Lipid binding pockets in S/x1 locked-1 (**a**) and locked-2 (**b**) conformations show clear lipid densities. Lipid binding pockets appear to be unoccupied in S/x2 locked-2 (**c**) and S/x1 closed (**d**-**e**) conformations showing absence of lipid density.

### Structural features of SARS-CoV-1 spikes in different conformations

To further understand conformational dynamics of SARS-CoV-1 spike, representative structures in each conformation category are summarized in **Fig. 3** with features highlighted (additional structures are shown **Fig. S5**). In SARS-CoV-1 S/x1 locked-1 protomers (**Fig. 3a**), a linoleic acid can be modelled in the density identified in a lipid binding pocket within the RBD. This pocket was previously identified in SARS-CoV-2 spike and it has been proposed to be a conserved feature in betacoronavirus spikes (Toelzer *et al*., 2020). By cryo-EM, we confirmed that the lipid is bound in a highly conserved fashion between SARS-CoV-1 and SARS-CoV-2 spikes. The aliphatic tail of the bound lipid is surrounded by aromatic amino acids lining the pocket. The pocket is gated by a small helix from a neighboring protomer to allow R395 within the gating helix to form salt bridge to the negatively charged lipid head group (**Fig. 3a, right panel**). Therefore, the bound lipid is interacting with RBDs of two neighboring protomers and these interactions are consistent to the proposal that they stabilize locked conformations (Toelzer *et al*., 2020). In the S/x1 locked-1 protomer, cryo-EM density for residues between 602 and 619 is not visible suggesting that these residues form a flexible loop (**Fig. 3a, left panel** and **Fig. 4a**). In an unusual dimer of locked-1 conformation SARS-CoV-2 spikes, formation of a large extended loop by this region was confirmed (Bangaru *et al*., 2020). In S/x1 locked-2 spike protomers (**Fig. 3b**), linoleic acid is bound in the RBD in an almost identical way as observed for the locked-1 protomer (**Fig. 3b, right panel**). Different from locked-1 spike protomers, residues between 602 and 619 in S/x1 locked-2 spike protomers refold into ordered structures showing clear cryo-EM density (**Fig. 3b, left panel** and **Fig. 4b**). The S/x2 locked-2 spike protomer displays cryo-EM density showing an ordered 602-619 region in Domain D (**Fig. 3c, left panel**), highly similar to S/x1 locked-2 spike protomer (**Fig. 3b, left panel**). Surprisingly, the lipid binding pocket is unoccupied even though the entrance to the pocket is closed by the gating helix (**Fig. 3c, right panel**). S/x3 spike was only observed to adopt a symmetrical locked-1 conformation, its protomers share very similar features as S/x1 locked-1 protomers, with bound lipid and residues 602-619 forming a flexible loop (**Fig. S5b**).

**Fig. 4.**
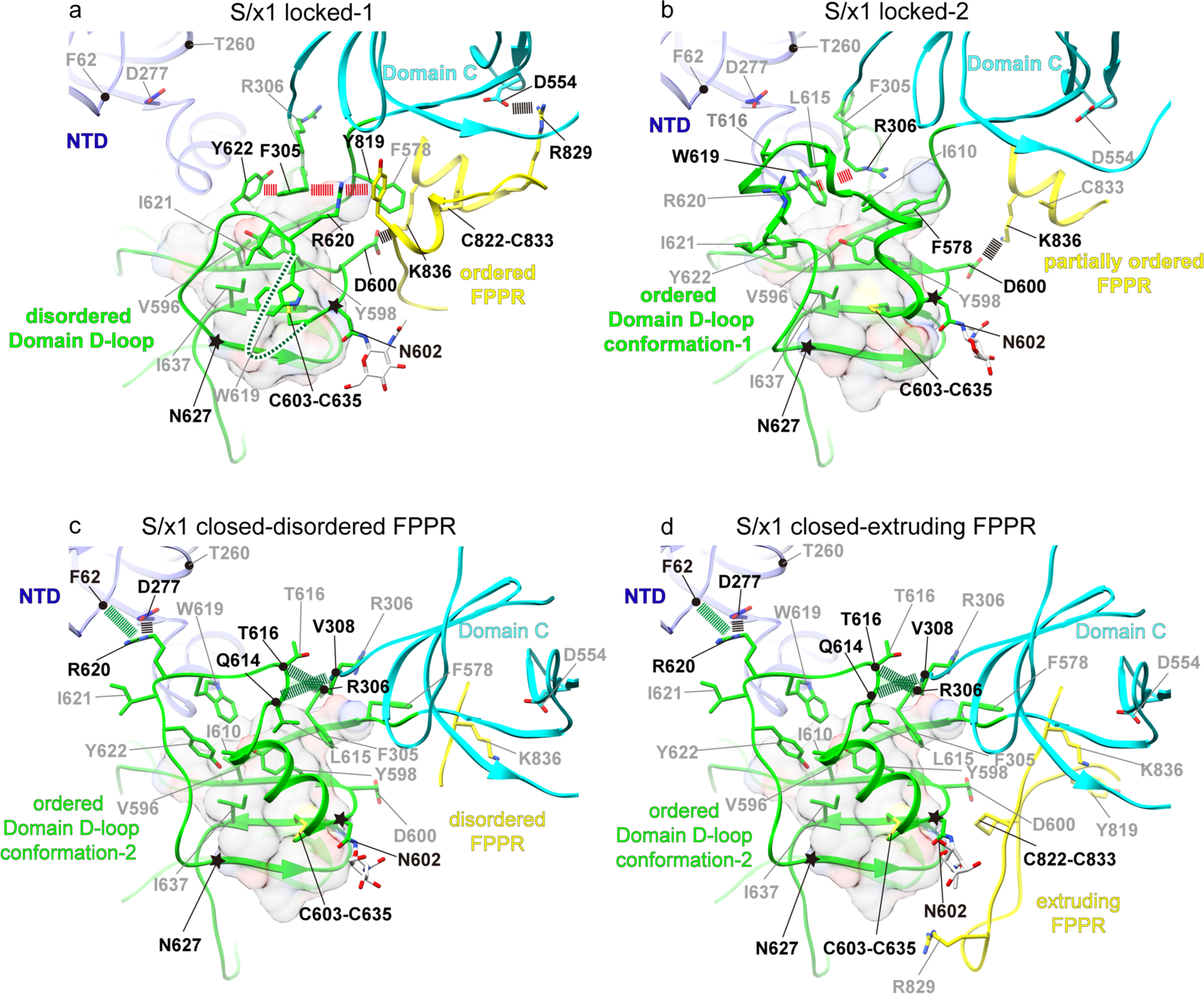
Structural rearrangements in Domain D among SARS-CoV-1 spikes of different conformations. **a**, Domain D and surrounding structural elements in locked-1 conformation. Residues N602-W619 within the Domain D-loop are disordered and represented by a green dashed line. Domain D-loop (residues N602-N627) where structural rearrangements occur between conformations are highlighted by star symbols. Cation-π and π-π interactions in the interdigitation interaction involving Y622, F305, R620 and Y819 are indicated by red dashed lines. Salt bridges are indicated by black dashed lines. Residues involved in featured interactions are highlighted by black labels. Hydrophobic residues (V596, Y598, and I637) within Domain D β-sheets are shown along with transparent molecular surface of Domain D hydrophobic core. A folded and ordered FPPR (yellow) is identified in locked-1 conformation. **b**, Domain D-loop in locked-2 conformation refolds to form ordered structures. A new cation-π interaction (red dashed line) is formed between R306 and W619. FPPR is partially disordered, the interaction between D600 of Domain D and K836 of FPPR is retained (black dashed line). **c-d**, Further refolding of Domain D-loop is observed in closed conformations and hydrophobic residues within the loop pack differently against the Domain D hydrophobic core. Interactions (black and green dashed lines) are formed to stabilize the folded Domain D-loop. **d**, FPPRs are normally disordered in spikes of closed conformation, an ordered extruding FPPR is observed for one protomer of closed S/x1 spike.

We identified clear densities consistent to biliverdin molecules in S/x1 locked-1, locked-2, locked-112 and locked-122 structures in a NTD pocket equivalent to the previously identified biliverdin binding pocket in SARS-CoV-2 S (Qu *et al*., 2022; Rosa *et al*, 2021) (**Fig. S5c**). Despite imaged under similar conditions, no density or very weak densities for biliverdin were observed in the biliverdin binding pockets of the other currently reported structures. In contrast, biliverdin densities were evident in spikes of locked and closed conformations for SARS-CoV-2 (Qu *et al*., 2022). We noticed certain degrees of sequence conservation within the largely hydrophobic biliverdin binding pocket between SARS-CoV-1 and SARS-CoV-2 S, but there are changes at several interacting residues (**Fig. S5c**). Molecular simulations suggested sequence changes in this pocket could affect biliverdin binding (Qu *et al*, 2021). It has been suggested that biliverdin binding could interfere with antibody binding (Cerutti *et al*, 2021; Rosa *et al*., 2021). Spike NTD has been previously observed to bind cofactors with unknown functions with regard to CoV life cycle, for example, folic acid has been identified to bind MERS-CoV spike NTD (Pallesen *et al*, 2017). SARS-CoV-1 and SARS-CoV-2 represent two distinct lineages of Sarbecoviruses (Guo *et al*, 2021), conservation of biliverdin binding pocket between their spikes, may suggest an unidentified physiological role for this molecule in Sarbecoviruses infection cycle.

Two distinct conformations were identified for protomers in the closed S/x1 asymmetrical spike (**Fig. 3d** and **e**). In both conformations, lipid binding pockets were unoccupied, consistent to the fact that the gating helices from neighboring protomers are dislocated leaving lipid binding pockets open (**Fig. 3d and e, right panels**). In both closed protomers, clear cryo-EM densities were observed for residues between 602 and 619 (**Fig. 3d and e, left panels and Fig. 4c** and **d**). However, they adopted a different structure comparing to the equivalent region in the locked-2 protomers (compared to **Fig. 3b** and **c, left panels**). FPPR has been found disordered in closed SARS-CoV-2 spikes, in line with this observation, FPPR is disordered in 2 of the 3 protomers within the SARS-CoV-1 S/x1 asymmetrical closed spike. However, surprisingly, one of the protomer has an FPPR forming an ordered extruding structure (**Fig. 3d, left panel)**. S/x2 spike can adopt a symmetrical closed conformation, spike protomer of this conformation has empty lipid binding pocket and disordered FPPR (**Fig. S5a**). Therefore, S/x2 closed protomer has almost identical features to the S/x1 closed protomer with a disordered FPPR (compare **Fig. S5a** and **Fig. 3e**).

### Domain D adopts different structures in SARS-CoV-1 spikes of different conformations

We summarized four representative structural arrangements around Domain D identified from determined cryo-EM structures. We examined interactions around Domain D in these structures in details. In locked-1 protomers, interdigitation interactions between Y622, F305, R620, and Y819 are formed through π-π and cation-π interactions (**Fig. 4a**). These interactions connect structural elements of Domain D (Y622, R620), Domain C-D hinge region (F305) and FPPR (F819) (**Fig. 4a**). Equivalent interdigitation interactions were observed in SARS-CoV-2 locked-1 spike. We have proposed that these interactions rigidify this area preventing structural transitions needed for RBD opening (Qu *et al*., 2022). Density for residues 602-619 is not visible and the equivalent region is known to form a large disordered loop in SARS-CoV-2 S adopting a locked-1 conformation (Bangaru *et al*., 2020) (**Fig. 4a**). Hydrophobic residues W619, I621 located at C-terminus of the loop interact with V596, Y598, and I637 of Domain D hydrophobic core (**Fig. 4a**). Both the interdigitation and hydrophobic interactions appear to hold the disordered loop in place. A salt bridge is formed between D600 of Domain D and K836 of FPPR (**Fig. 4a**). D600 is equivalent to D614 in SARS-CoV-2 spike and D614G is the earliest mutation became fixed in SARS-CoV-2 spike since the pandemic and all subsequent variants bear this mutation (Grubaugh *et al*, 2020; Korber *et al*, 2020). Likely stabilized by the participated interactions, FPPR is ordered in locked-1 conformation consisting of two small helices connected by a disulfide between C822-C833 (**Fig. 4a**). Presence of this structural motif is known to be incompatible with RBD in “up” position (Xiong *et al*., 2020).

In the locked-2 conformation, the highly dynamic region between residues 602-627 in Domain D (Domain D-loop) undergoes drastic refolding (**Fig. 4b**) to adopt a fully ordered “Domain D-loop conformation-1” with several mini-helices within the loop. Interdigitation interactions connecting Domain D, Domain C-D hinge region and FPPR in the locked-1 conformation are broken. Residues between 602 and 619 become ordered in the locked-2 conformation (**Fig. 4b**). A change of Domain C-D hinge region structure allows residues R306 and F578 to interact with W619 and Domain D hydrophobic core by cation-π and hydrophobic interactions, respectively. These interactions likely still rigidify this region to maintain the more tightly packed locked-2 conformation (**Fig. S6** and **Fig. S7**). Most interactions between FPPR and other parts of spike, including the interdigitation interactions are lost, only the salt-bridge between K836 of FPPR and D600 of Domain D is retained. FPPR is partially ordered with clear density only for the helix formed by residues 830-838 (**Fig. 4b** and **Fig. 3b, c**).

Further structural changes around Domain D occur in the S/x1 closed conformations, a large structural change caused bending and rotation of Domain C-D hinge region allowing residue F305 to interact with the Domain D hydrophobic core (**Fig. 4c and d**). The large Domain D-loop (residues 602-627) assumes a different ordered conformation (here we refer it as “Domain D-loop conformation-2”), the three mini-helices in Domain D-loop of locked-2 conformation unfold and a longer helix between 602-610 is formed (**Fig. 4c** and **d**). This conformational change allowed R620 to interact with NTD F62 mainchain carbonyl and D277 sidechain carboxylate by hydrogen bond and salt bridge respectively (**Fig. 4c** and **d**). Due to this refolding, W619 and Y622 interact with the Domain D hydrophobic core differently (**Fig. 4c** and **d**). While the above described interactions involve residues conserved between SARS-CoV-1 and SARS-CoV-2 spikes, cryo-EM density was not observed for Domain D-loop in SARS-CoV-2 spike of closed conformation, suggesting that the equivalent Domain D-loop adopts a different flexible conformation (Qu *et al*., 2022). We identified numerous residues in or around Domain D differ between SARS-CoV-1 and SARS-CoV-2 spikes (**Fig. S8**) making it difficult to pin down which amino acid changes result in different Domain D structural dynamics between the two spikes. We noted though, if T260 in SARS-CoV-1 S is substituted for an arginine found in the equivalent position of SARS-CoV-2 S (**Fig. 4c-d** and **Fig. S8**), it would be in close proximity to R620 of Domain D-loop, repulsion between the two positively charged residues would be unfavorable for Domain D-loop to adopt the ordered “Domain D-loop conformation-2” that is featured in closed conformations of SARS-CoV-1 spike.

### SARS-CoV-1 and SARS-CoV-2 spikes sample similar conformations

Previously we found domain movement in a protomer alters trimer packing resulting in differences in spike quaternary structure between locked and closed conformations (Qu *et al*., 2022). To further understand conformational dynamics of SARS-CoV-1 spike, S1 of SARS-CoV-1 spikes of different conformations were structurally aligned using Domain D as the reference (**Fig.5** and **Fig. S9**). The overlaps show that drastic refolding of Domain D between locked-1 and locked-2 conformations alters the structure of Domain C-D hinge region and affects the structure of FPPR (**Fig. 5a**). Despite these extensive structural changes, positioning of NTD and RBD is largely unaffected, only small shifts in NTD and RBD positions are observed (**Fig. 5a**). Similarly, the two closed conformations are almost superimposable when aligned, despite structural differences in FPPR (**Fig. 5b**). In contrast, comparison of SARS-CoV-1 S locked-1 and locked-2 conformations to closed conformations revealed that positioning of RBD differ considerably between locked and closed conformations (**Fig. 5c and d**). The overlap of locked and closed conformations revealed that RBD movement originates from the bendings in the Domain C-D hinge region (**Fig. 5c and d**). We previously also identified similar characteristics for SARS-CoV-2 S, structural transition from locked conformations to closed conformation is associated with refolding of Domain D, bending of Domain C-D hinge region, and movement of RBD (Qu *et al*., 2022). Overlaps of SARS-CoV-1 and SARS-CoV-2 S1 of different conformations reveal that spikes of the two CoVs share very similar locked-1, locked-2 and closed conformations (**Fig. 5e-g**).

**Fig. 5.**
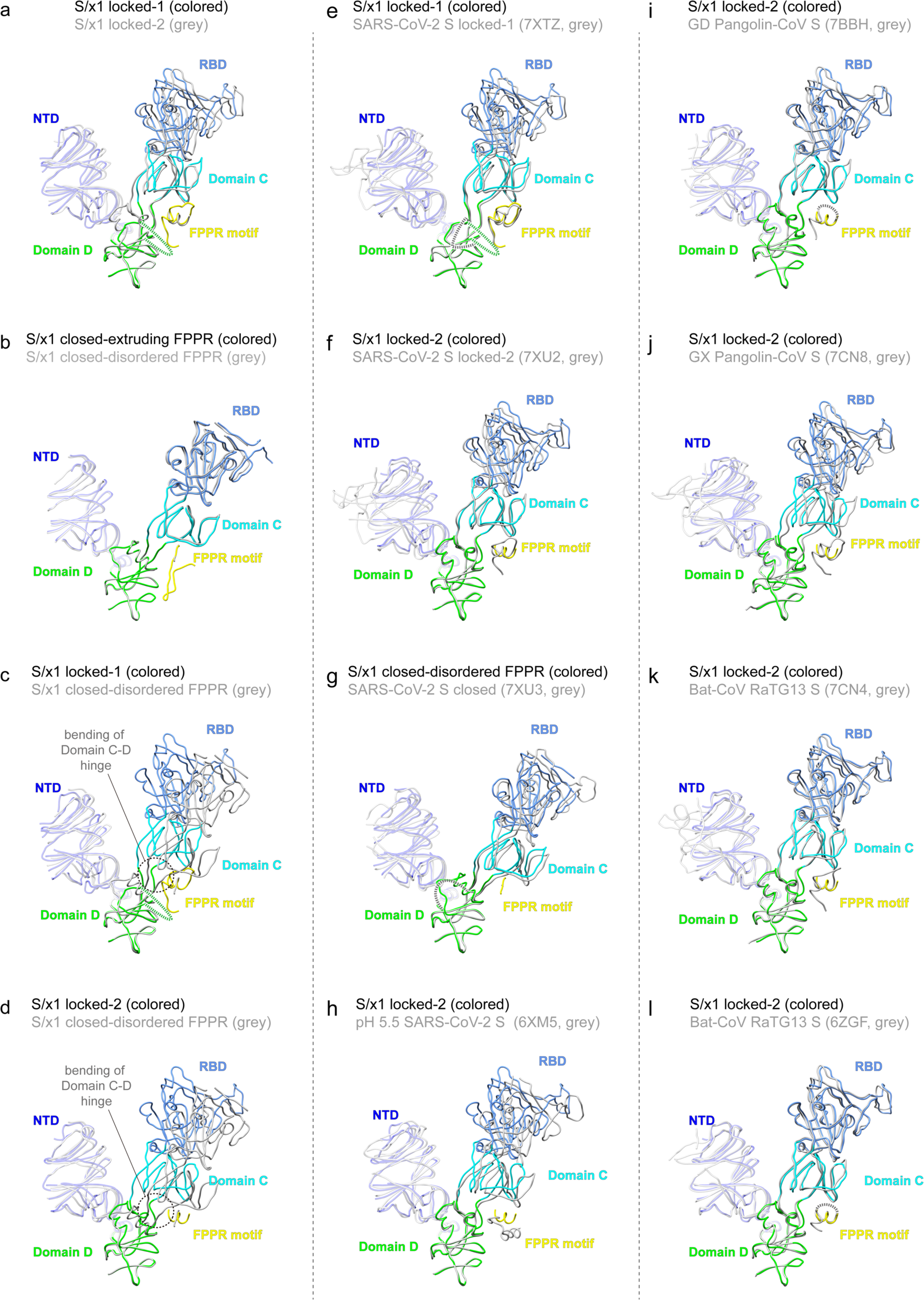
SARS-CoV-1 and SARS-CoV-2 spikes are highly dynamic adopting multiple shared conformations, other SARSr-CoV spikes exhibit reduced dynamics. **a-d,** Comparison of SARS-CoV-1 spike S1 portions of different conformations. Structures are shown in ribbon representations. Positions of S1 structural domains remain largely unchanged between locked-1 and locked-2 conformations despite structural rearrangements in Domain D. **b,** An overlap of two S/x1 closed conformations reveals highly similar positions of S1 structural domains. **c-d,** Overlaps of locked-1 and locked-2 conformations to closed conformation reveal a large motion of RBD between locked and closed conformations originating from the bendings in Domain C-D hinge region. **e-f,** Overlaps of SARS-CoV-1 and SARS-CoV-2 spike S1 structures of related conformations to show similarities (PDBs: 7XTZ, 7XU2, 7XU3 (Qu *et al*., 2022); 6XM5 (Zhou *et al*., 2020)). **i-l,** Overlaps of SARS-CoV-1 and SARSr-CoV spike S1 structures of related conformations to identify similarities (PDBs: 7BBH (Wrobel *et al*., 2021); 7CN8, 7CN4 (Zhang *et al*., 2021b); 6ZGF (Wrobel *et al*., 2020)).

Several details are different: first, whilst FPPR is fully ordered in SARS-CoV-2 S of both locked-1 and locked-2 conformations (**Fig. 5e and f**), FPPR is only fully ordered in SARS-CoV-1 S of locked-1 conformation (**Fig. 5e**). Second, SARS-CoV-2 S in closed conformation is characterized by its disordered Domain D-loop and FPPR (**Fig. 5g**), whereas in SARS-CoV-1 S, Domain D-loop is refolded to adopt an ordered structure but distinct from that in locked-2 conformation (**Fig. 5g and Fig. 4b, c, d**). Third, we identified an asymmetrical closed conformation for SARS-CoV-1 S in which FPPR is dynamic adopting either extruding loop or disordered structures, whereas FPPR is usually fully disordered in closed conformation SARS-CoV-2 S (**Fig. 5b**). Previously, dynamics was observed for FPPR when SARS-CoV-2 spike was treated with acidic pH (**Fig. 5h**) (Zhou *et al*, 2020).

## Discussion

Class I viral membrane fusion glycoproteins are evolved to have metastable prefusion conformation (Harrison, 2008). Introduction of disulfides into viral fusogenic glycoproteins has been proven useful to stabilize flu HA (Godley *et al*, 1992; Lee *et al*, 2015), HIV GP160 (Sanders *et al*, 2013) and paramyxovirus F proteins (McLellan *et al*, 2013; Wong *et al*, 2016) in various prefusion conformations for mechanistic, structural and vaccinology studies. Here, we introduced x1, x2, x3 disulfides previously designed for SARS-CoV-2 S into SARS-CoV-1 S, this allowed structural characterization of previously unobserved conformations for SARS-CoV-1 spike. Newly determined structures revealed that SARS-CoV-1 S is able to form symmetrical locked-1, locked-2 trimers or asymmetrical trimers made up by locked-1 and locked-2 protomers in various combinations. An asymmetrical closed trimer was also observed with one of the protomer featuring an extruding FPPR. Therefore, together with previously determined structures, it is confirmed that SARS-CoV-1 spike can adopt locked-1, locked-2, closed and RBD “up” open conformations. SARS-CoV-1 and SARS-CoV-2 spikes appear to sample similar conformations and they both exhibit complex conformational dynamics.

Structures of a few other Sarbecovirus spikes have also been determined. SARSr-CoV spikes, namely Guangdong (GD) pangolin-CoV S (Wrobel *et al*, 2021), Guangxi (GX) pangolin-CoV S (Zhang *et al*, 2021b), and RaTG13 bat-CoV (Wrobel *et al*., 2020; Zhang *et al*., 2021b) S, only adopt the locked-2 conformation, when imaged by cryo-EM (**Fig. 5i-l**). Therefore, SARSr-CoV spikes are different from SARS-CoV-1 and SARS-CoV-2 spikes exhibiting reduced structural dynamics. Alternatively, closed/open conformations of SARSr-CoV spikes may not be stable enough to withstand cryo-EM sample preparation procedures and that the more structurally rigid spikes in locked conformations were preferentially imaged.

For SARS-CoV-2 (Toelzer *et al*., 2020) and SARSr-CoV spikes (Zhang *et al*., 2021b), locked conformations are often associated with lipid binding in RBD and lipid binding has been proposed to stabilize locked spike trimers (Toelzer *et al*., 2020). In this study, association of lipid binding and locked conformations was observed for SARS-CoV-1 S/x1 and S/x3 spikes. Previously, we and others found that low-pH was able to convert closed/open SARS-CoV-2 spike to locked conformations (Qu *et al*., 2022; Zhou *et al*., 2020). Based on these observations and the egress pathway characterized for betacoronaviruses (Ghosh *et al*, 2020), we have proposed that locked conformations are associated with virus particle assembly in low-pH, lipid rich endomembrane structures (Qu *et al*., 2022).

In this study we confirmed that locked-1 and locked-2 conformations are shared between SARS-CoV-1 and SARS-CoV-2 spikes and locked-2 conformations are shared among all characterized Sarbecovirus spikes. Structural characterization reveals that, although details differ between locked-1 and locked-2 conformations, restraining RBD movement by interactions between Domain D and Domain C-D hinge region is shared by locked conformations of Sarbecovirus spikes. Conservation of RBD restraining mechanisms in locked conformations among Sarbecovirus spikes suggests locked conformations may play a conserved function in Sarbecovirus life cycle.

Previously, only closed and open conformations were observed for purified SARS-CoV-1 S (**Fig. S9g-l**). We have attempted to convert native SARS-CoV-1 S ectodomain (S/native) without engineered disulfide to locked conformation by prolonged incubation under low-pH (pH 5.5 for 24 hr), however, we only observed open and closed spikes after incubation (**Fig. S4**). In line with previously determined SARS-CoV-1 S structures (Gui *et al*., 2017; Yuan *et al*., 2017), Domain D-loops in closed and open spike structures determined under low-pH are ordered, adopting “Domain D-loop conformation-2”, showing almost identical conformations as previously detailed for the closed conformation (**Fig. S9** and **Fig. S10**). Of note, during preparation of this manuscript, a preprint reported a subpopulation of purified insect cell expressed (presumably expressed in low-pH insect cell media) SARS-CoV-1 spike ectodomain adopted a locked-2 conformation with lipid bound in RBD (Toelzer *et al*, 2022). We speculate that unlike SARS-CoV-2 S, where Domain D-loops are disordered in closed and open conformations, breaking interactions involving the ordered Domain D-loops in closed and open conformations represents a barrier to revert closed/open spike to locked conformations for SARS-CoV-1 S. This difference reflects how changes in Domain D may affect dynamics of Sarbecovirus spikes. Indeed, D614G mutation (Yurkovetskiy *et al*, 2020; Zhang *et al*, 2021a) and proteolytic cleavage by cathepsin L (Zhao *et al*, 2022), both within Domain D, are known to affect dynamics of SARS-CoV-2 spike.

In summary, SARS-CoV-1 and SARS-CoV-2 spikes exhibit complex conformational dynamics among Sarbecovirus spikes. Biological functions of these different conformations remain to be further confirmed or elucidated. In addition, underlying mechanisms by which different Sarbecovirus spikes sample various conformations differently and the associated biological consequences remain poorly understood. Future “*in-situ*” structural studies under more conditions may provide further clues for these questions. Nevertheless, in this study, we demonstrated that introduction of stabilizing disulfides in SARS-CoV-1 spike is useful for capturing otherwise transient conformations allowing more thorough characterization of spike dynamics. We further demonstrated engineered S/x3 spike as an immunogen. Together with previously determined SARS-CoV-2 spike structures, we believe conserved features characterized for SARS-CoV-1 and SARS-CoV-2 spikes should help further understanding of Sarbecovirus spike function.

**Table 1.** Cryo-EM data collection, refinement and validation statistics

|  | S/x1 locked-1<br>(EMDB-34417)<br>(PDB: 8H0X) | S/x1 locked-112<br>(EMDB-34418)<br>(PDB: 8H0Y) | S/x1 locked-122<br>(EMDB-34419)<br>(PDB: 8H0Z) | S/x1 locked-2<br>(EMDB-34420)<br>(PDB: 8H10) | S/x1 closed<br>(EMDB-34421)<br>(PDB: 8H11) |
| --- | --- | --- | --- | --- | --- |
| **Data collection and processing** |  |  |  |  |  |
| Magnification |  |  | 81000 |  |  |
| Voltage (kV) |  |  | 300 |  |  |
| Electron exposure (e<sup>-</sup>/Å<sup>2</sup>) |  |  | 50 |  |  |
| Defocus range (μm) |  |  | -0.8~-2.0 |  |  |
| Pixel size (Å) |  |  | 1.095 |  |  |
| Symmetry imposed | C3 | C1 | C1 | C3 | C1 |
| Initial particle images (no.) | 4604239 | 4604239 | 4604239 | 4604239 | 4604239 |
| Final particle images (no.) | 58285 | 91477 | 70911 | 22532 | 504118 |
| Map resolution (Å) | 2.57 | 2.85 | 2.99 | 2.99 | 2.72 |
| FSC threshold | 0.143 | 0.143 | 0.143 | 0.143 | 0.143 |
| Map resolution range (Å) | 2.46-5.18 | 2.74-4.74 | 2.87-4.65 | 2.85-4.74 | 2.55-4.92 |
| **Refinement** |  |  |  |  |  |
| Initial model used (PDB code) | 7XTZ | 7XTZ/7XU2 | 7XTZ/7XU2 | 7XU2 | 5X58 |
| Model resolution (Å) | 2.7 | 2.9 | 3.0 | 3.1 | 2.8 |
| FSC threshold | 0.5 | 0.5 | 0.5 | 0.5 | 0.5 |
| Map sharpening *B* factor (Å<sup>2</sup>) | -26 | -30 | -29 | -34 | -48 |
| **Model composition** |  |  |  |  |  |
| Non-hydrogen atoms | 24804 | 24940 | 24797 | 24882 | 23477 |
| Protein residues | 3078 | 3086 | 3078 | 3078 | 2932 |
| Ligands | 777 | 833 | 776 | 861 | 560 |
| ***B* factors (Å<sup>2</sup>)** |  |  |  |  |  |
| Protein | 46.20 | 64.68 | 65.79 | 54.21 | 38.19 |
| Ligand | 61.70 | 80.90 | 81.68 | 78.21 | 74.64 |
| **R.m.s. deviations** |  |  |  |  |  |
| Bond lengths (Å) | 0.004 | 0.003 | 0.006 | 0.007 | 0.004 |
| Bond angles (°) | 0.988 | 0.73 | 0.725 | 1.054 | 0.665 |
| **Validation** |  |  |  |  |  |
| MolProbity score | 1.19 | 1.10 | 1.20 | 1.20 | 1.04 |
| Clashscore | 3.31 | 2.14 | 4.17 | 2.98 | 2.50 |
| Poor rotamers (%) | 0.26 | 0.3 | 0.3 | 0.34 | 0 |
| **Ramachandran plot** |  |  |  |  |  |
| Favored (%) | 97.69 | 97.46 | 97.98 | 97.43 | 97.97 |
| Allowed (%) | 2.31 | 2.54 | 2.02 | 2.57 | 2.03 |
| Disallowed (%) | 0 | 0 | 0 | 0 | 0 |

|  | S/x2 closed<br>(EMDB-34422)<br>(PDB: 8H12) | S/x2 locked-2<br>(EMDB-34423)<br>(PDB: 8H13) | S/x3 locked-1<br>(EMDB-34424)<br>(PDB: 8H14) |
| --- | --- | --- | --- |
| **Data collection and processing** |  |  |  |
| Magnification |  | 45000x |  |
| Voltage (kV) |  | 200 |  |
| Electron exposure (e<sup>-</sup>/Å<sup>2</sup>) |  | 62 |  |
| Defocus range (μm) |  | -0.8~-2.4 |  |
| Pixel size (Å) |  | 0.88 |  |
| Symmetry imposed | C3 | C3 | C3 |
| Initial particle images (no.) | 967643 | 967643 | 82693 |
| Final particle images (no.) | 82882 | 192457 | 22958 |
| Map resolution (Å) | 3.96 | 3.37 | 3.28 |
| FSC threshold | 0.143 | 0.143 | 0.143 |
| Map resolution range (Å) | 3.79~9.19 | 3.26~6.92 | 3.11~10.00 |
| **Refinement** |  |  |  |
| Initial model used (PDB code) | 5x58 | 7XU2 | 7XTZ |
| Model resolution (Å) | 4.04 | 3.45 | 3.38 |
| FSC threshold | 0.5 | 0.5 | 0.5 |
| Map sharpening B factor (Å<sup>2</sup>) | -137 | -105 | -39.5 |
| Model composition |  |  |  |
| Non-hydrogen atoms | 22233 | 24330 | 24762 |
| Protein residues | 2793 | 3036 | 3075 |
| Ligands | 378 | 630 | 777 |
| B factors (Å<sup>2</sup>) |  |  |  |
| Protein | 43.73 | 27.86 | 25.88 |
| Ligand | 86.83 | 55.30 | 46.88 |
| R.m.s. deviations |  |  |  |
| Bond lengths (Å) | 0.004 | 0.004 | 0.003 |
| Bond angles (°) | 0.758 | 1.096 | 0.597 |
| **Validation** |  |  |  |
| MolProbity score | 1.58 | 1.21 | 1.05 |
| Clashscore | 3.92 | 2.27 | 2.03 |
| Poor rotamers (%) | 0.08 | 1.03 | 0 |
| Ramachandran plot |  |  |  |
| Favored (%) | 93.91 | 96.79 | 97.63 |
| Allowed (%) | 6.09 | 3.21 | 2.37 |
| Disallowed (%) | 0 | 0 | 0 |

|  | S/native low-pH closed<br>(EMDB-34425)<br>(PDB: 8H15) | S/native low-pH open<br>(EMDB-34426)<br>(PDB: 8H16) |
| --- | --- | --- |
| **Data collection and processing** |  |  |
| Magnification |  | 45000x |
| Voltage (kV) |  | 200 |
| Electron exposure (e<sup>-</sup>/Å<sup>2</sup>) |  | 60 |
| Defocus range (μm) |  | -0.8~-2.5 |
| Pixel size (Å) |  | 0.88 |
| Symmetry imposed | C3 | C1 |
| Initial particle images (no.) | 1375445 | 1375445 |
| Final particle images (no.) | 201293 | 308387 |
| Map resolution (Å) | 3.07 | 3.28 |
| FSC threshold | 0.143 | 0.143 |
| Map resolution range (Å) | 2.95~9.13 | 3.07~9.02 |
| **Refinement** |  |  |
| Initial model used (PDB code) | 5X58 | 5X5B |
| Model resolution (Å) | 3.07 | 3.28 |
| FSC threshold | 0.5 | 0.5 |
| Map sharpening *B* factor (Å<sup>2</sup>) | -68.5 | -63.5 |
| Model composition |  |  |
| Non-hydrogen atoms | 22571 | 20534 |
| Protein residues | 2890 | 2630 |
| Ligands | 392 | 0 |
| *B* factors (Å<sup>2</sup>) |  |  |
| Protein | 22.64 | 27.80 |
| Ligand | 49.45 | 0 |
| R.m.s. deviations |  |  |
| Bond lengths (Å) | 0.003 | 0.004 |
| Bond angles (°) | 0.72 | 0.82 |
| **Validation** |  |  |
| MolProbity score | 1.22 | 1.45 |
| Clashscore | 3.11 | 4.44 |
| Poor rotamers (%) | 0.12 | 0.44 |
| Ramachandran plot |  |  |
| Favored (%) | 97.41 | 96.47 |
| Allowed (%) | 2.59 | 3.53 |
| Disallowed (%) | 0 | 0 |

## Author contributions

X.X conceived the study; X.Z. expressed and purified spikes for cryo-EM and other experiments based protein production protocols developed by X.X.; X.Z. performed negative stain EM with assistance from B.L., Q.C., X.Zhong and L.J.; X.Z. performed spike reduction experiments with assistance from Q.C.; Y.Z. and Q.W. performed animal experiments under the supervision of L.C. and J.Z.; Y.Z. performed pseudovirus neutralization assays under the supervision of J.Z.; X.Z. prepared cryo-EM grids with assistance from Y.L. and J.W.; Y.L., L.F., X.Z. and J.W. collected cryo-EM data under the supervision of P.W. and J.H.; Z.L. and X.Z. processed cryo-EM data under the supervision of X.X.; X.Z., Z.L. and X.X. built structure models; X.Z. and X.X. analyzed cryo-EM structures; X.Z and Z.L. prepared figures with assistance from X.X.; X.X., X.Z. wrote the paper with input from all authors.

## Acknowledgements

We thank Katarzyna Ciazynska, Andrew Carter (MRC-LMB, Cambridge, UK) and John Briggs (Max Planck Institute of Biochemistry, Martinsried, Germany) for initial design of constructs and development of protein production protocols. We thank Kun Qu (Yong Loo Lin School of Medicine, National University of Singapore) for assistance in cryo-EM data processing. This study was supported by the Guangdong-Hong Kong-Macau Joint Laboratory of Respiratory Infectious Diseases (2019B121205010 to JH) and Natural Science Fund of Guangdong Province (2021A1515011289 to XX), National Natural Science Foundation of China (2021000192 to XX). XX acknowledges start-up grants from the Chinese Academy of Sciences and Bioland Laboratory (GRMH-GL).

---

**Fig. S1.**
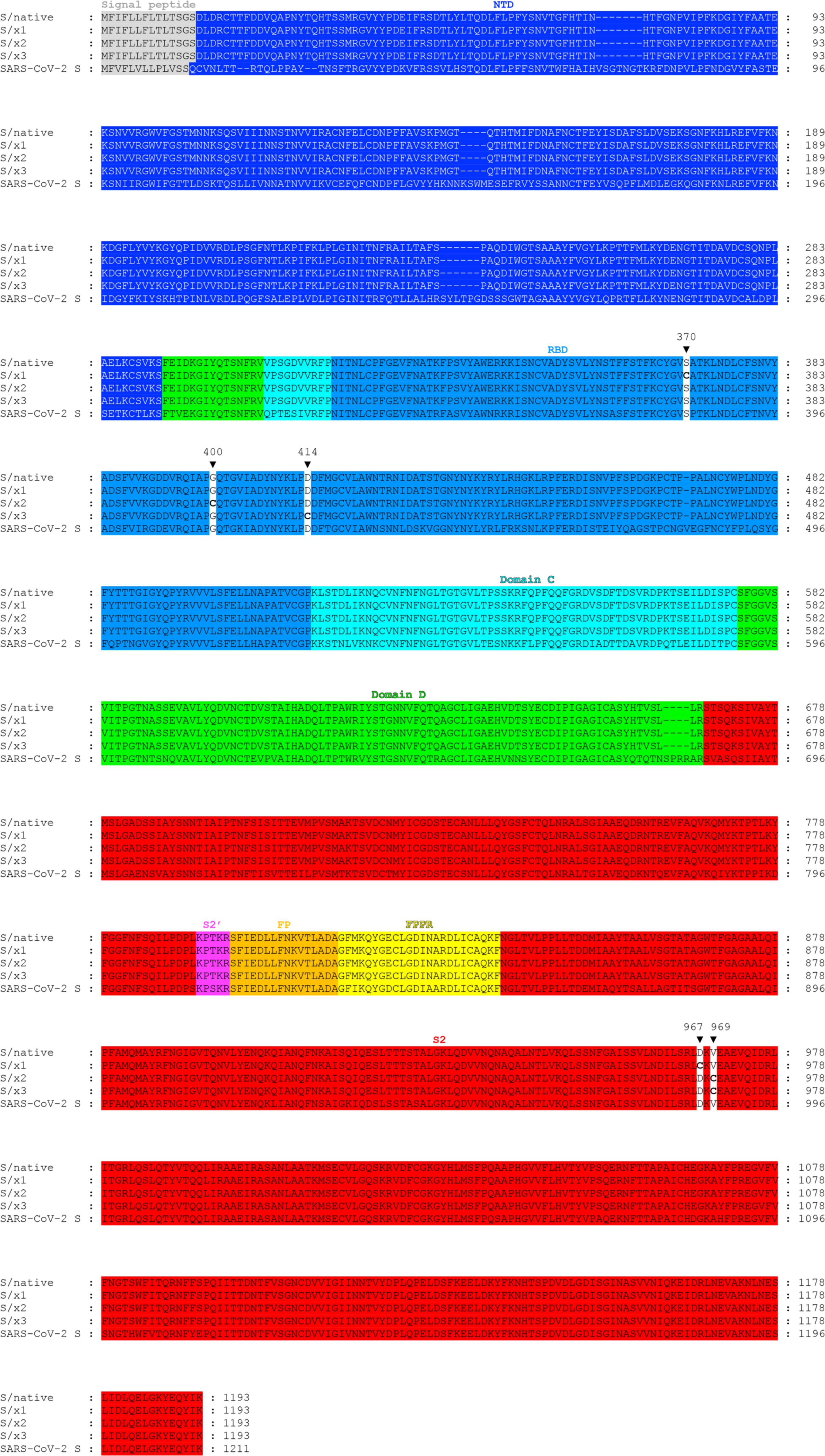
Sequence alignment of S/native, S/x1, S/x2, S/x3 and SARS-CoV-2 S. Signal peptide is colored in grey. S1 domains - NTD, RBD, Domain C and Domain D are highlighted in blue, light-blue, cyan and green respectively. S2 is colored in red. S2 structural elements - S2’, fusion peptide (FP), and fusion peptide proximal region (FPPR) are highlighted magenta, orange and yellow respectively. Black triangles mark positions at which cysteines were introduced and indicate amino acid residue numbers according to SARS-CoV-1 S protein sequence. Introduced cysteines are shown in bold.

**Fig. S2.**
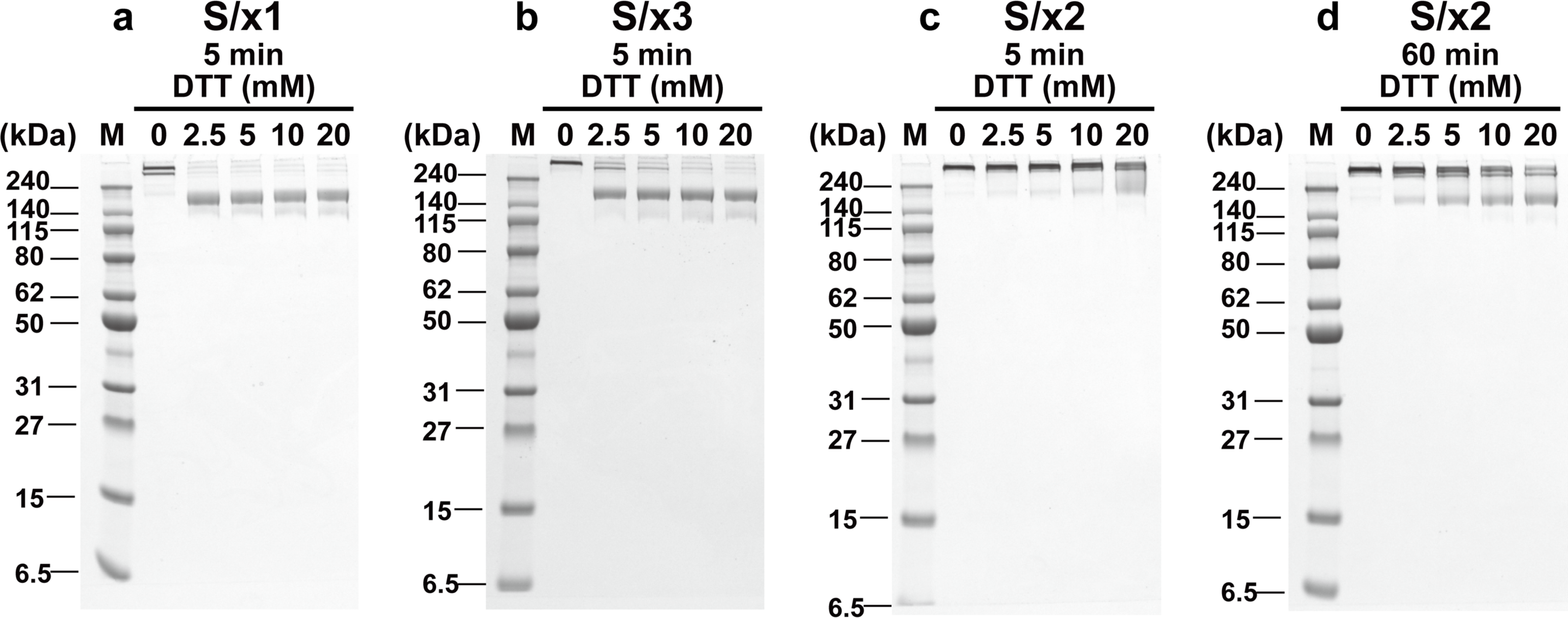
Engineered disulfides in S/x1, S/x2 and S/x3 spike proteins show different sensitivities to reduction by DTT. **a-c**, Reduction of engineered disulfides was tested by incubating S/x1, S/x3 and S/x2 spike proteins with indicated concentrations of DTT for 5 min under native conditions. DTT was quenched by excess amount of iodoacetamide before SDS-PAGE. **c**, Incomplete reduction of S/x2 spike was observed after 5 min DTT incubation. **d**, S/x2 spike proteins was incubated with indicated concentrations of DTT for 60 minutes under native conditions before iodoacetamide quenching and SDS-PAGE.

**Fig. S3.**
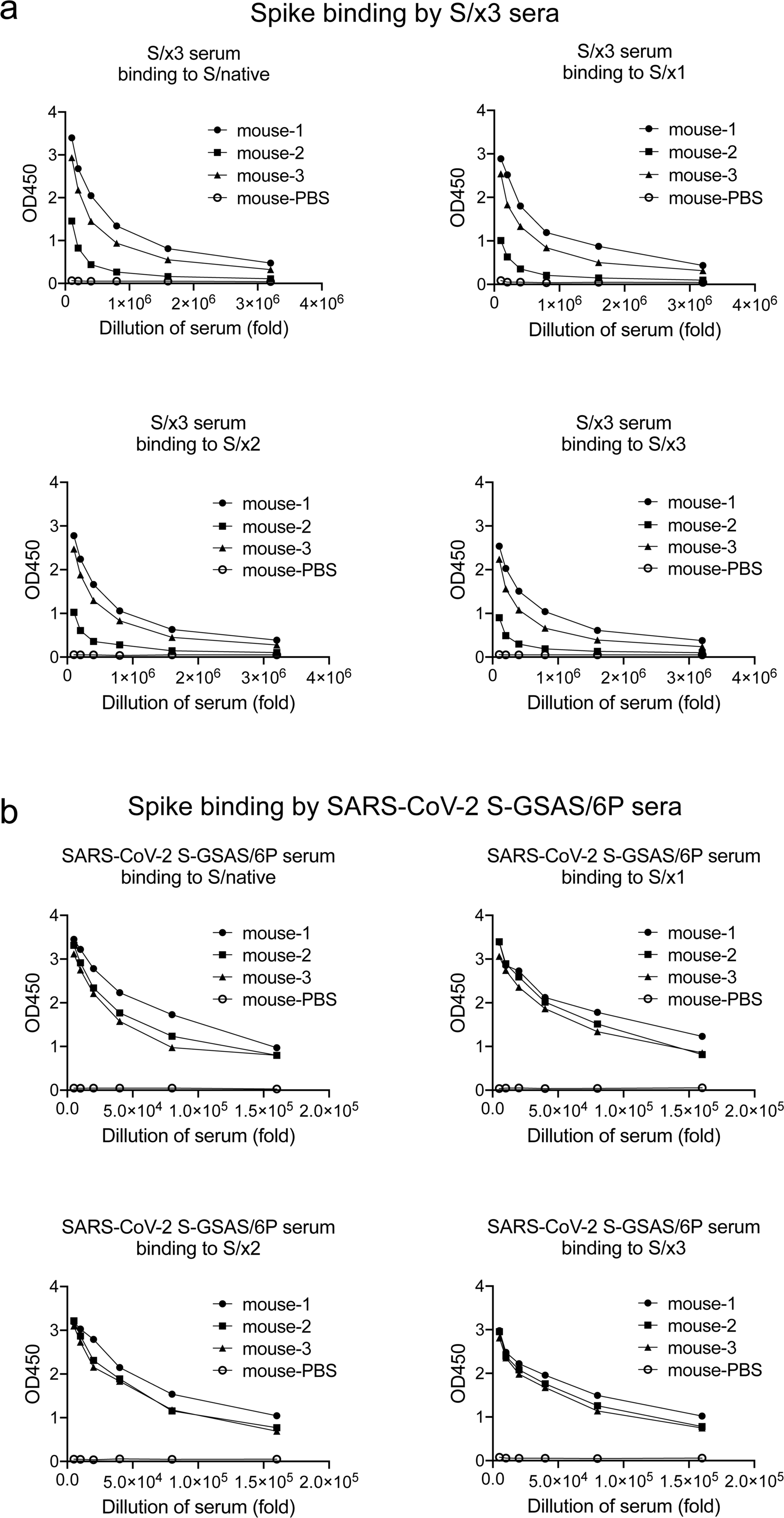
Binding of SARS-CoV-1 S/native/, S/x1, S/x2 and S/x3 spike proteins by mouse immune sera. **a-b**, mouse immune sera were raised by priming and boosting with purified SARS-CoV-1 S/x3 or SARS-CoV-2 S-GSAS/6P spike (10 μg protein for each dose). Binding to different spike proteins were assessed by ELISA. Titration curves were generated by dilution of immune sera.

**Fig. S4.**
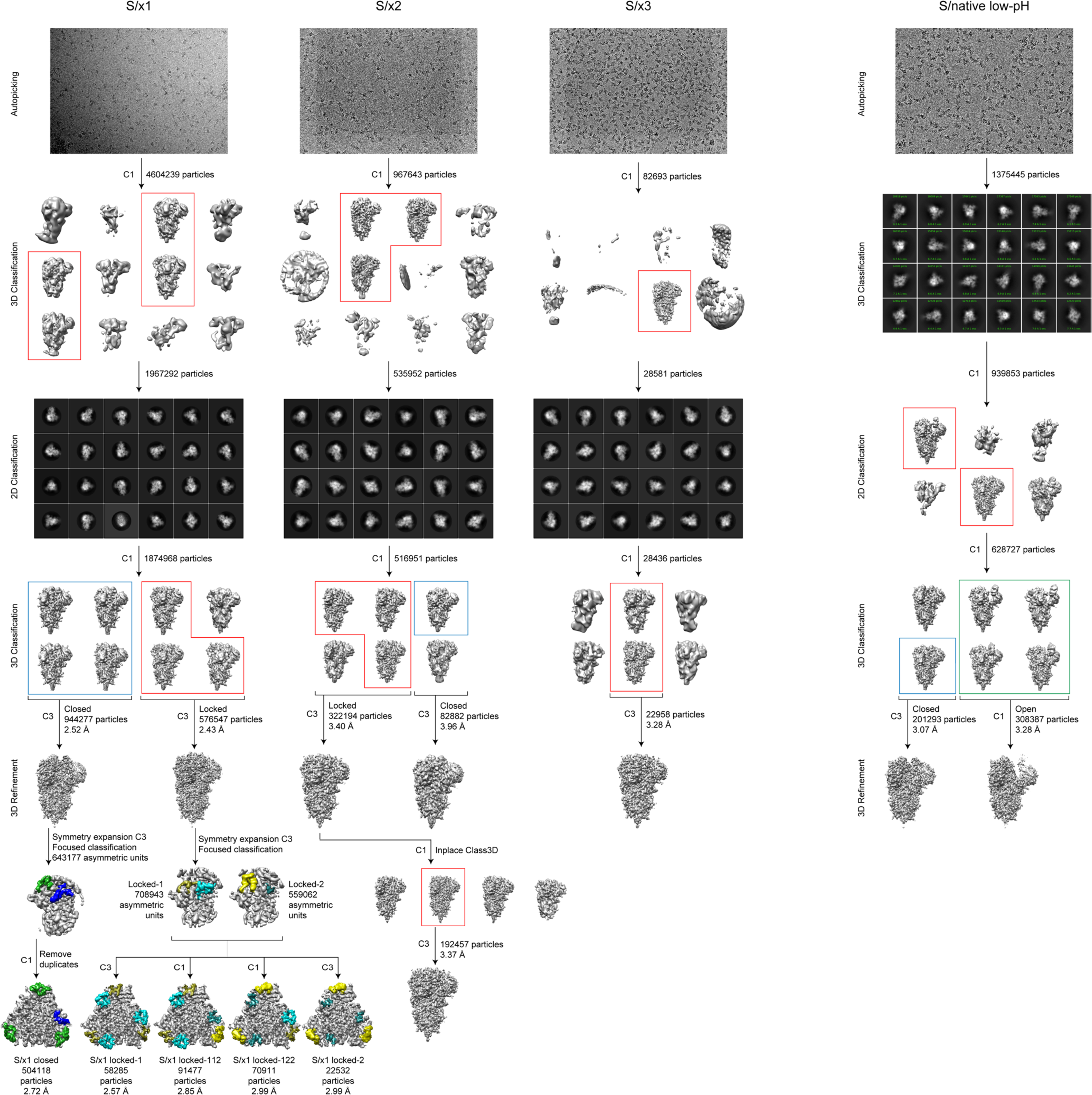
Cryo-EM processing flow-chart for reported structures. Data processing pipelines are illustrated for cryo-EM datasets collected for SARS-CoV-1 S/x1, S/x2, S/x3 and S/native (under low-pH) spike proteins. After automated particle picking, 3D and 2D classification steps were used to remove contaminating objects. Further 3D classification procedures were used to identify structures with distinct features.

**Fig. S5.**
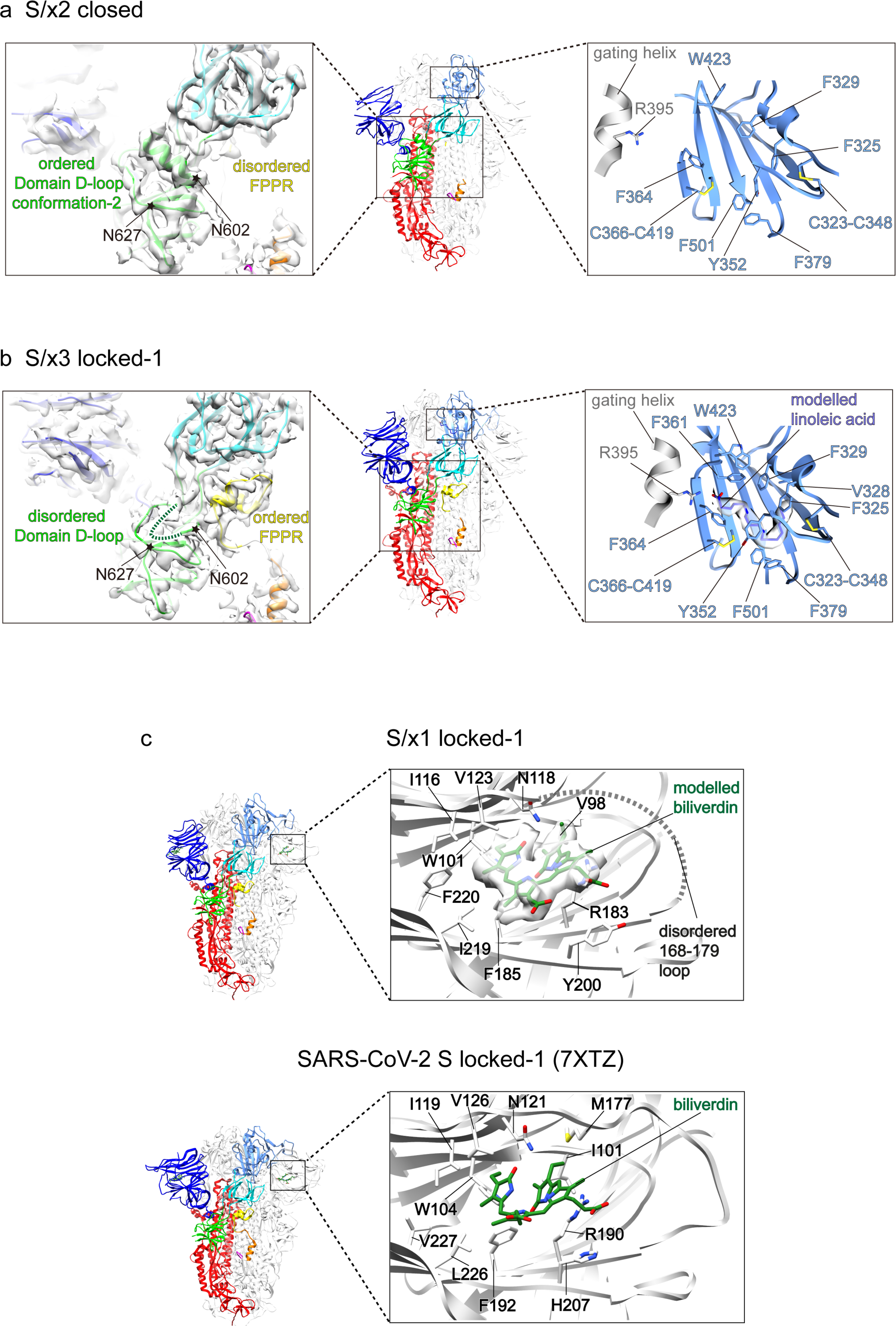
Structural features in S/x2 closed, S/x3 locked-1 spike structures and putative biliverdin binding pocket in SARS-CoV-1 S. **a-b,** Middle panels show molecular models for S/x2 closed and S/x3 locked-1 spike structures, domains and structural elements are colored for the featured protomers as in **Fig. 2**. Left panels show cryo-EM densities of Domain D and surrounding regions with fitted molecular models. Regions between N602-N627 where structural rearrangements occur are highlighted by star symbols. Right panels show lipid binding pockets within the two spike RBDs. Hydrophobic amino acid sidechains forming the lipid binding pocket are shown in stick representations. Gating helices from the neighboring RBDs are shown in grey, the head group interacting R395 are shown in stick representations. **c,** Top panel, putative biliverdin binding site identified in SARS-CoV-1 S/x1 of locked-1 conformation. Putative biliverdin density is shown in surface representation, modelled biliverdin is shown in green, sidechains of interacting residues are shown in stick representations. The disordered loop between residues 168-179 in the vicinity of biliverdin binding site is represented by the dashed line. Bottom panel, the previously identified biliverdin binding site in SARS-CoV-2 S of locked-1 conformation (PDB: 7XTZ) (Qu *et al*., 2022), biliverdin and equivalent interacting residues are shown in stick representations. Many biliverdin interacting residues show conservation between SARS-CoV-1 and SARS-CoV-2 spikes.

**Fig. S6.**
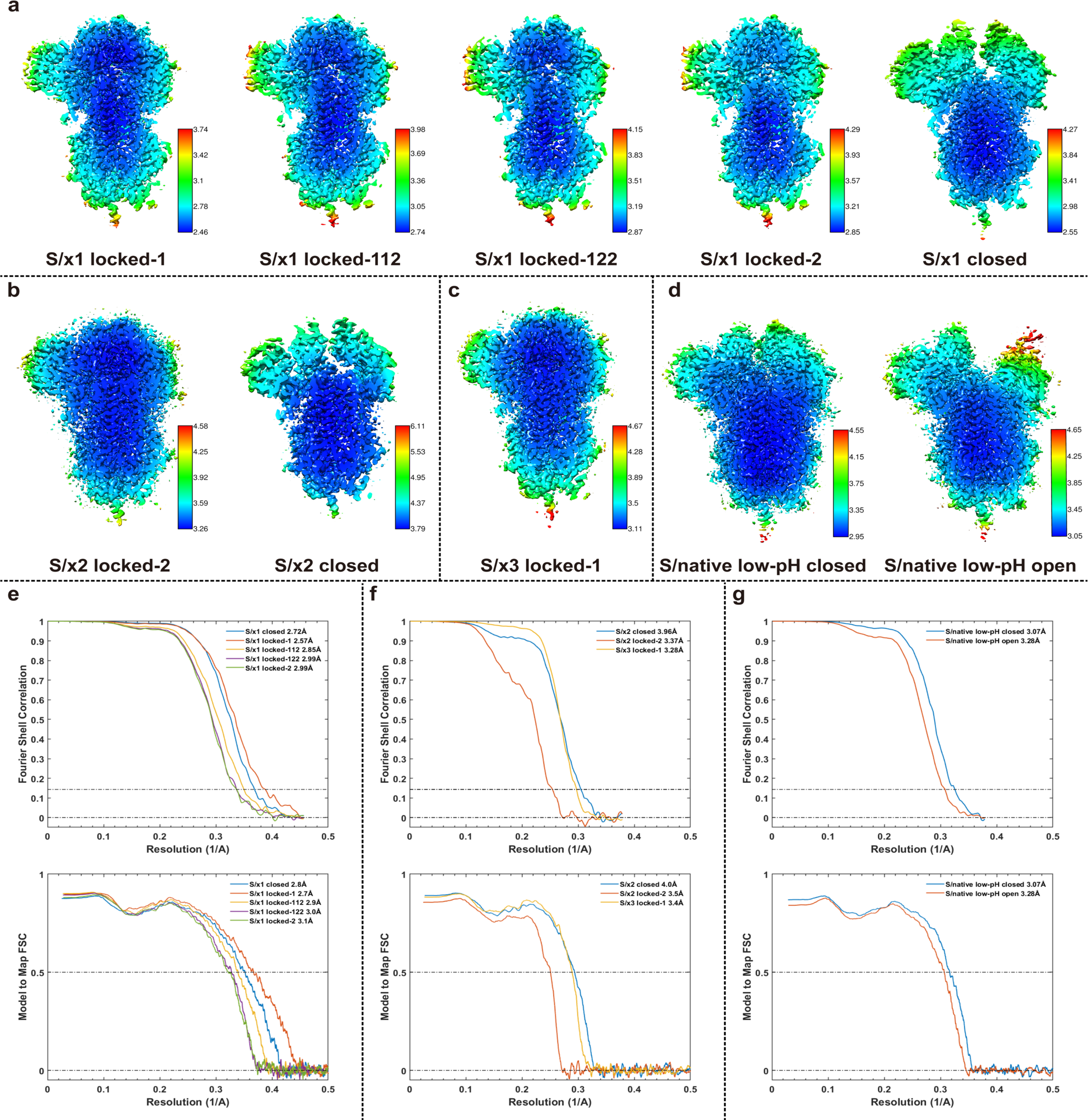
Resolution assessment of cryo-EM structures. **a-d**, Local resolution maps for S/x1, S/x2, S/x3 and low-pH S/native cryo-EM structures. **e-g**, Fourier shell correlations for S/x1, S/x2, S/x3 and low-pH S/native cryo-EM structures. Top panel, global resolution assessment by Fourier shell correlation at the 0.143 criterion. Bottom panel, correlations of model vs map by Fourier shell correlation (FSC) at the 0.5 criterion.

**Fig. S7.**
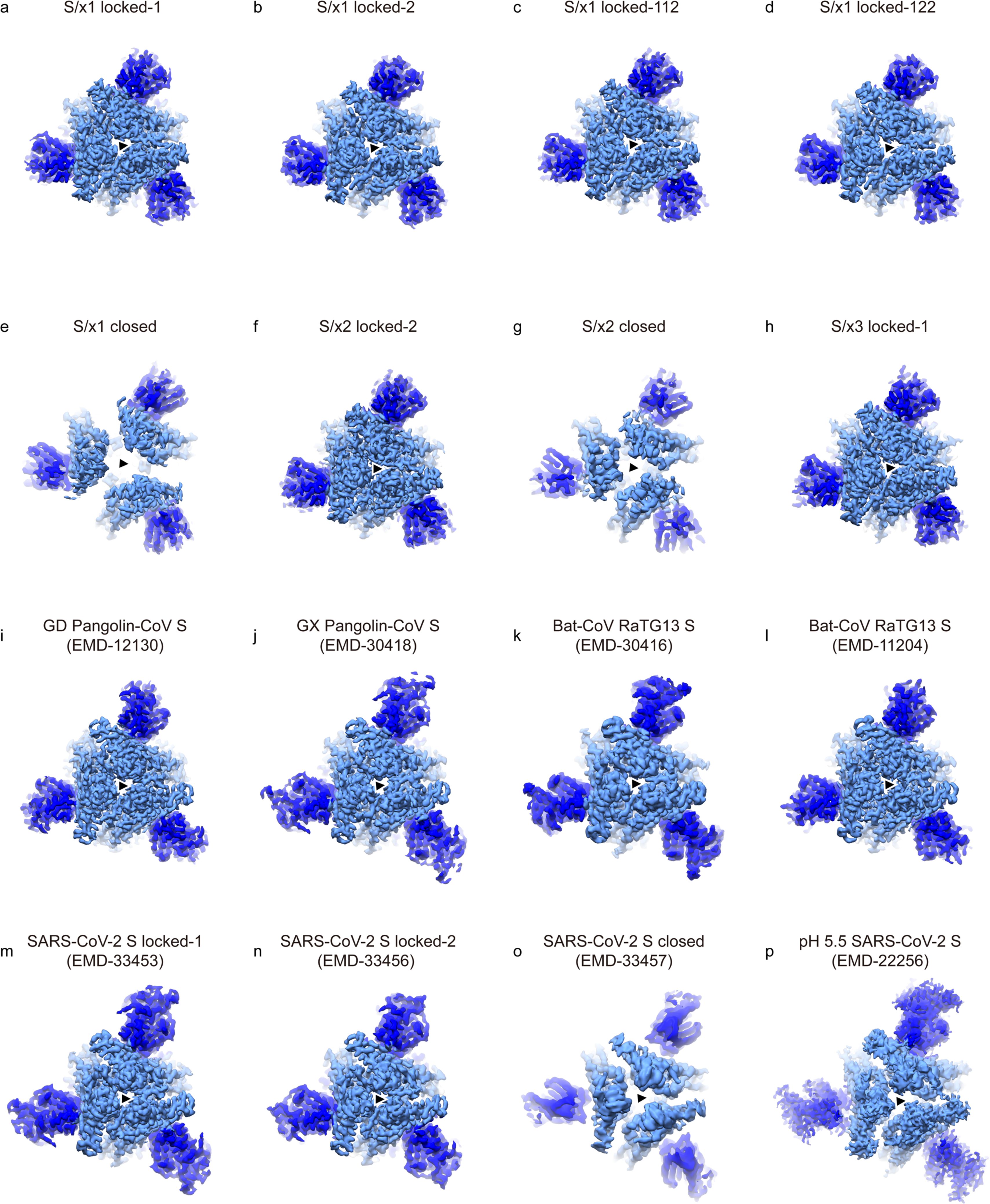
Locked and closed spikes show difference in trimer packing. **a-p**, Top views of cryo-EM densities showing Sarbecovirus spikes adopting different conformations. NTD and RBD are colored in blue and light blue respectively. The three-fold axes are indicated by black triangles. Locked SARS-CoV-1 spikes (**a-d**; **f** and **h**), animal SARSr-CoV spikes (**i**, EMD-12130 (Wrobel *et al*., 2021); **j**, EMD-30418 (Zhang *et al*., 2021b); **k**, EMD30416 (Zhang *et al*., 2021b); **l**, EMD-11204 (Wrobel *et al*., 2020)) and locked SARS-CoV-2 spikes (**m**, EMD-33453 (Qu *et al*., 2022); **n**, EMD-33456 (Qu *et al*., 2022)) show more compact packing of RBDs comparing to closed spike trimers (**e**; **g**; **o,** EMD-33457 (Qu *et al*., 2022)). **p**, The structure of SARS-CoV-2 S determined under pH 5.5 (EMD-22256 (Zhou *et al*., 2020)) exhibits trimer packing intermediate of closed and locked conformations.

**Fig. S8.**
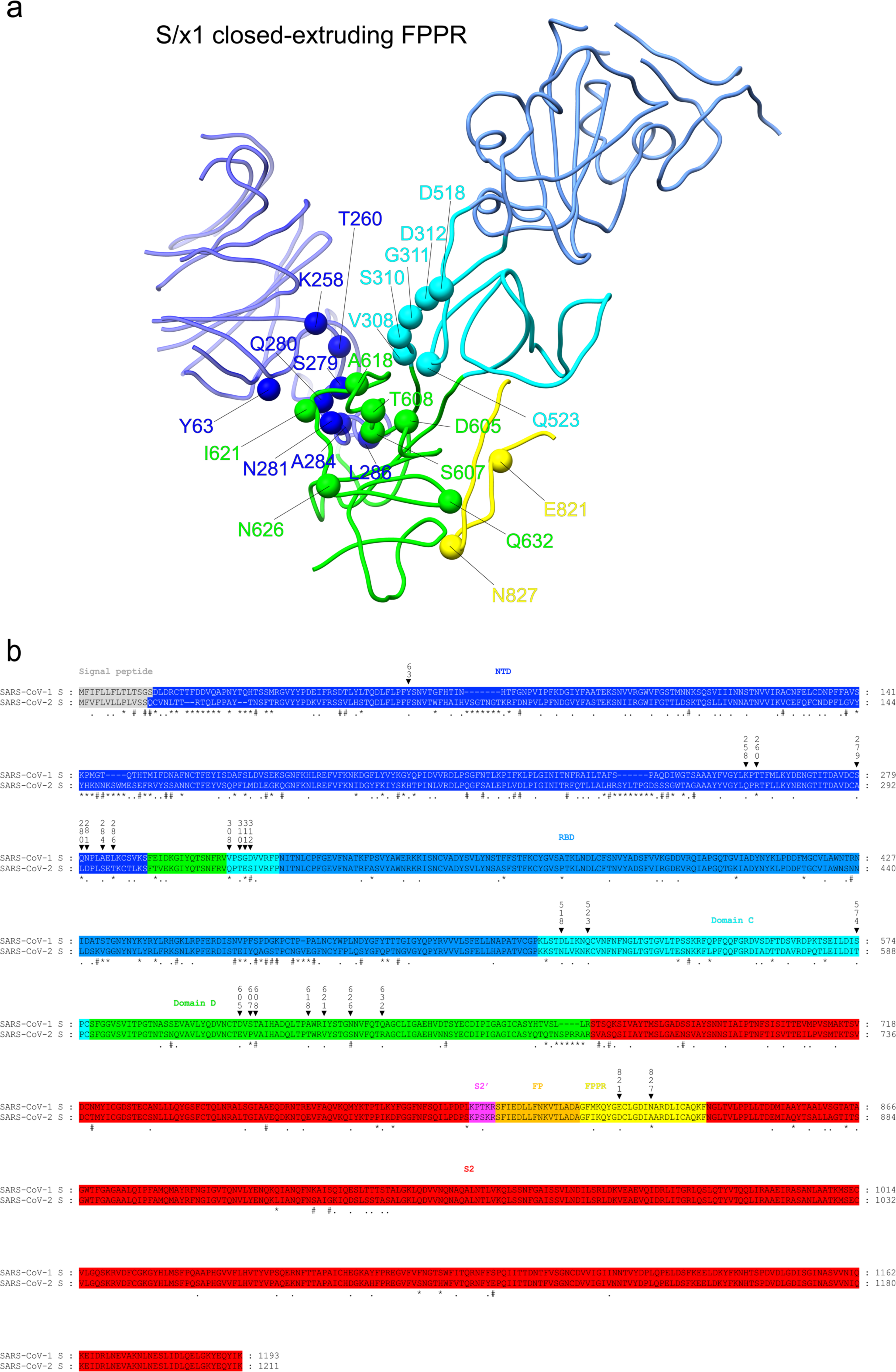
Amino acid differences between SARS-CoV-1 and SARS-CoV-2 in and around Domain D. **a**, SARS-CoV-1 S/x1 closed conformation structure with extruding FPPR is shown in a ribbon representation. Spheres indicate amino acid differences between SARS-CoV-1 and SARS-CoV-2 S sequences in/around Domain D. **b**, Alignment of SARS-CoV-1 and SARS-CoV-2 S ectodomain sequences. Structural domains and elements are shaded in indicated colors. Asterisk (*), hash (#), and period (.) above amino acid sequences indicate amino acids differ without similar properties, with weakly similar properties, and with strongly similar properties respectively. Triangles above amino acid sequences indicate amino acid differences shown in (**a**) with their amino acid positions in SARS-CoV-1 S sequence numbered.

**Fig. S9.**
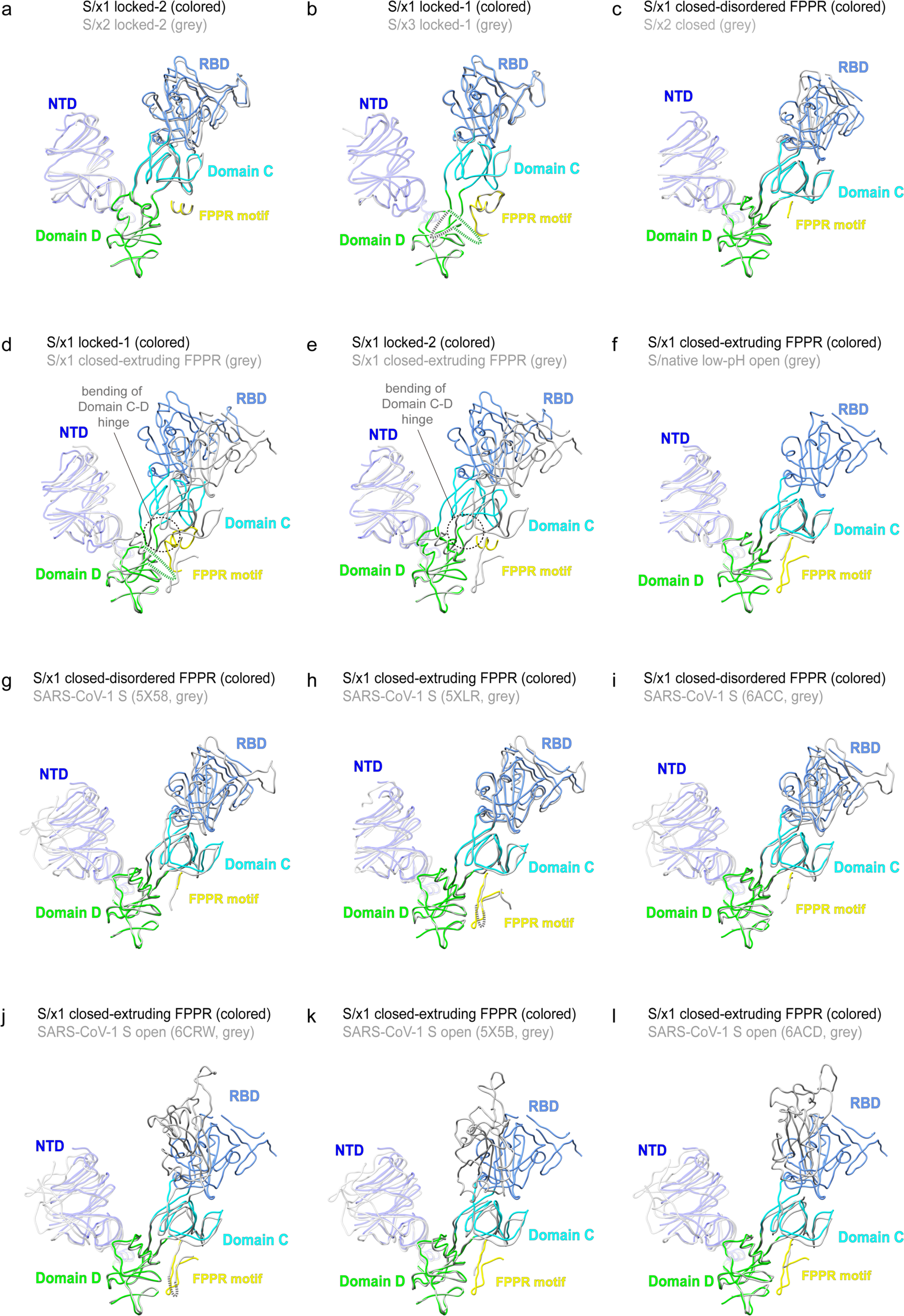
Superpositions of S1 portions of selected SARS-CoV-1 spike structures. **a-e**, Superpositions of S1 structures of S/x1, S/x2, S/x3 and low-pH S/native spikes of different conformations. **g-l**, Superpositions of S1 structure of S/x1 closed spike with extruding FPPR to available SARS-CoV-1 S structures in the PDB (PDBs: 5X58, 5X5B (Yuan *et al*., 2017); 5XLR (Gui *et al*., 2017); 6ACC, 6ACD (Song *et al*, 2018); 6CRW (Kirchdoerfer *et al*, 2018)). NTD, RBD, Domain C, Domain D and FPPR are colored in blue, ligh blue, cyan, green and yellow respectively. Missing loops are indicated by dashed lines.

**Fig. S10.**
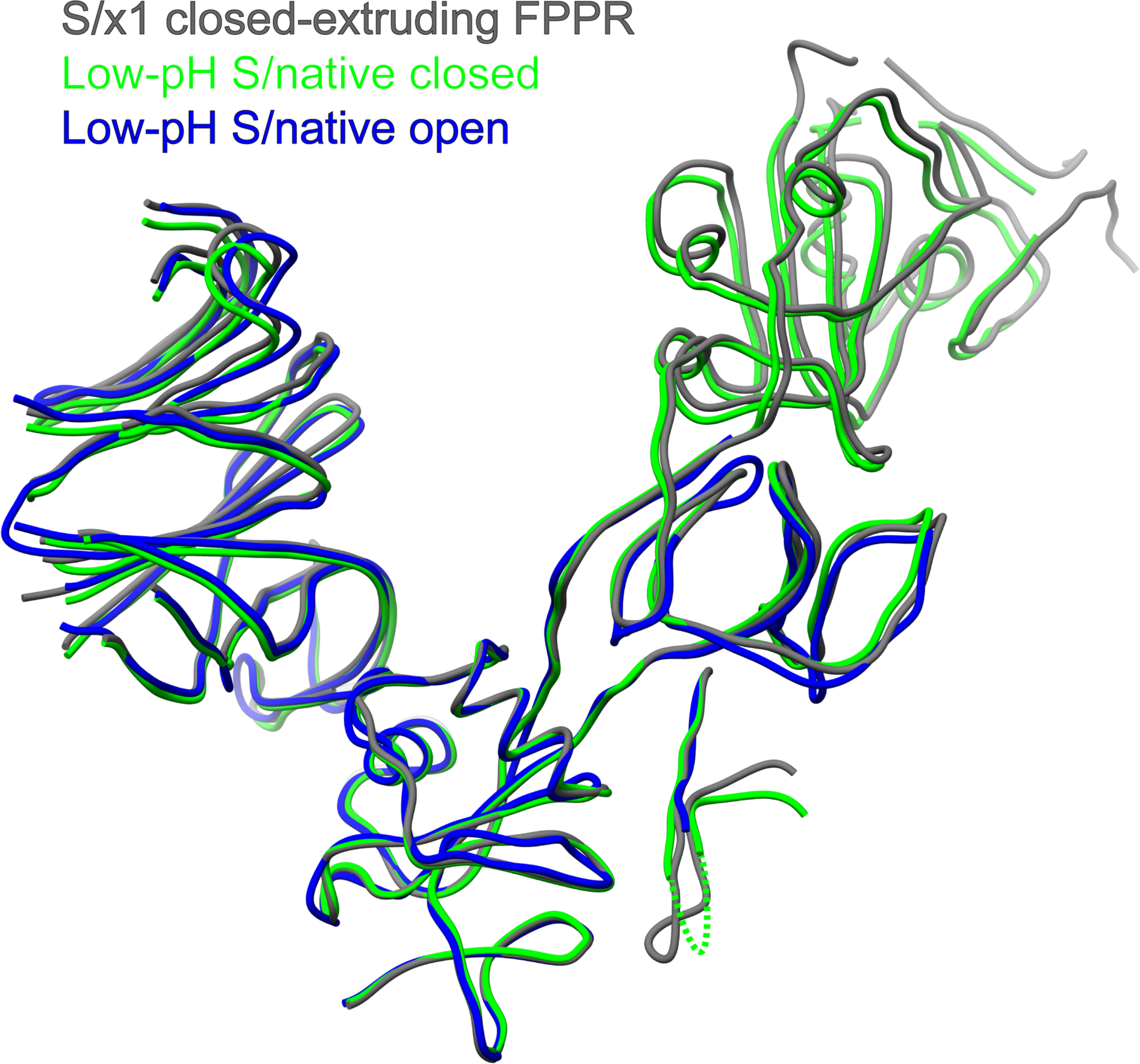
Comparison of S1 structures of S/x1 closed conformation and S/native closed and open conformations.

## Materials and Methods

### Protein expression and purification

S/native construct was generated by cloning S gene of SARS-CoV-1 (SARS coronavirus Tor2, NCBI:txid227984) between amino acid residues 15-1193 into a modified pcDNA3.1 vector previously described (Walls *et al*, 2019; Xiong *et al*., 2020). The S/native construct encodes an exogenous N-terminal signal peptide MGILPSPGMPALLSLVSLLSVLLMGCVAETGT derived from µ-phosphatase and a C-terminal extension GSGR*ENLYFQ*GGGGSGYIPEAPRDGQAYVRKDGEWVLLSTFLGHHHHHH. The C-terminal extension contains a tobacco etch virus (TEV) protease cleavage site (italic), a T4 trimerization foldon (underlined) and a hexa-Histidine-tag (bold). This construct is called S/native. S/x1 (S370C and D967C), S/x2 (G400C and V969C) and S/x3 (D414C and V969C) containing the introduced cysteines were generated by PCR.

Spike proteins were expressed by transient transfection in Expi293 cells, 1 mg of DNA was transfected into 1 L of cells using PEI transfection reagent. After transfection, cells were cultured at 33 °C for 5 days. Supernatants were harvested by centrifugation and 25 mM phosphate pH 8.0, 5 mM imidazole, 300 mM NaCl were supplemented. Supernatant was recirculated onto a 5-ml Talon Cobalt column for ∼2.5 times and the column was washed by 100 ml buffer A (25 mM phosphate pH 8.0, 5 mM imidazole, 300 mM NaCl). Spike protein was eluted by a linear gradient using increasing concentration of Buffer B (25 mM phosphate pH 8.0, 500 mM imidazole, 300 mM NaCl). Purified spike proteins were quality checked by SDS-PAGE and buffered exchanged into PBS and frozen at -80 °C until further use.

### Negative-staining EM

Spike proteins were diluted to 0.06 mg/ml and 3µl samples were applied onto glow-discharged (15 mA, 45 s glow) carbon-coated copper grids. Protein samples were absorbed for 1 minute, before grids were washed with water once. Samples were stained twice with 0.75% (w/v) uranyl formate solution. Grids were air dried and imaged in an FEI Tecnai G2 Spirit transmission electron microscope operating at 120 keV equipped with a CCD camera.

### Disulfide bond reduction under native conditions

S/x1, S/x2, and S/x3 spike proteins were incubated with 0, 2.5, 5, 10, and 20 mM DTT for 5 min or 60 min at room temperature in PBS. Reactions were quenched by addition of 55 mM iodoacetic acid (IAA) for 10 min in the dark at room temperature. Reaction mixtures were mixed with 4× non-reducing loading buffer and were analysed by SDS-PAGE.

### Animal Immunization

6-to 8-week-old BALB/c females were immunized intramuscularly with 10 μg of purified protein mixed with 100 μg of Sigma Adjuvant System (SAS) in a total volume of 100 μl of PBS. After a 4-week interval, mice were boosted with the same antigen in the same formulation. Day 28 after the priming immunization and day 7 after boost immunization, sera were collected from immunized mice to detect the antigen-specific humoral response by ELISA and pseudovirus neutralization assay.

### Pseudovirus neutralization assay

Pseudotyped lentiviruses were produced in 293T cells as previously described (Feng *et al*, 2020) by co-transfecting a plasmid expressing SARS-CoV-1 S protein, a packaging vector and a reporter vector carrying an expression cassette of firefly luciferase. The 4× serially diluted heat-inactivated (HI)-serum (56 ℃ for 30 min) were incubated with the SARS-CoV-1 pseudotyped virus at 37 °C for 1 h. The mixture was subsequently incubated with HeLa-ACE2 cells for 72h. The cells were washed twice with PBS and lysed with lysis buffer before measuring luciferase activity. The neutralization titers were calculated as antibodies dilutions at which the luciferase activity was reduced to 50% of that from the virus-only wells.

### ELISA assay to determine antibody binding activity to SARS-CoV-1 spikes

96 well assay plates were coated with 100 µl per well of SARS-CoV-1 S/native, S/x1, S/x2, or S/x3 spike proteins at 1 µg/ml in PBS overnight at 4 °C. After standard washing, 10% fetal calf serum (FBS, 200 µl per well) was added to block for 2 hours at 37 °C. After washing wells three times with PBS-0.1% Tween 20 (MP Biomedicals), 100 µl semilogarithmic dilutions of antisera in PBS were added to each well and incubated for 2 hours at 37°C. After washing wells 3 times with PBS-0.1% Tween 20, plates were incubated with 1:10000 dilutions of HRP-labelled Goat Anti-Mouse IgG (H + L) (Jackson ImmunoResearch Laboratories) in 10% FBS for 1 h at 37°C. After washing wells 6 times with PBS-0.1% Tween 20, 100 µl per well of TMB (3,3’,5,5’-tetramethylbenzidine) solution (Merck Millipore) was added and developed for 10 min at room temperature. Reactions were stopped by adding 50 µl 2 M sulphuric acid and OD values at 450 nm were measured in a plate reader.

### Cryo-EM sample preparation

Immediately before cryo-EM sample preparation, spike samples in PBS (S/x1 at 4.7 mg/ml, S/x2 at 6.5 mg/ml, S/x3 at 2.4 mg/ml, S/native at 6.68 mg/ml) were thawed and TEV protease (2.5 mg/ml in PBS) was added to a 10:1 spike/TEV protease w/w ratio. The mixtures were incubated at room temperature for 1 h before used for grids preparation. For the low-pH S/native spike sample, after the TEV protease incubation, per 9 µl of the protein solution was incubated with 1 µl of 1 M pH 5.5 sodium acetate buffer (giving final pH of 5.54 for the protein solution) overnight at room temperature. Quantifoil R1.2/1.3 copper grids were glow-discharged (15 mA, 30 s) before used for specimen preparation. 3 µl protein solution was mixed with 0.3 µl 1% octyl-glucoside (final concentrations of OG were 0.1%) immediately before applied to grid. Grids were blotted (3s, force 4) and rapidly plunge-frozen in liquid ethane using a Vitrobot (Thermo Fisher Scientific) in 100% humidity at 22 ℃. Grids were screen in a Talos Arctica G2 electron microscope (Thermo Fisher Scientific) operated by EM facility of Guangzhou Institutes of Biomedicine and Health. Suitable grids with clear side- and top-views of spikes were identified and datasets were collected in the Talos Arctica G2 electron microscope, or a Titan Krios G3i electron microscope (Thermo Fisher Scientific) operated by EM center of Southern University of Science and Technology (SUSTech), Shenzhen.

### Cryo-EM data collection

The Talos Arctica electron microscope was operating at 200 keV and Serial EM software v3.8.7 was used for data collection. On a K3 direct detection camera (Gatan), micrographs were acquired at a nominal magnification of 45,000×, with a calibrated pixel size of 0.88 Å and a defocus range of -0.8 to -2.5 μm. For the S/x2 and S/x3 samples, each movie was exposed for 1.6 s with a dose rate of 30 e^-^/pixel/second fractionated into 27 frames, resulting a total dose of 62 e^-^/Å^2^. For the low-pH S/native sample, each movie was exposed for 1.8 s with a dose rate of 26 e^-^/pixel/second fractionated into 27 frames, resulting a total dose of 60 e^-^/Å^2^.

The Titan Krios G3i electron microscope was operating at 300 keV and EPU software (Thermo Fisher Scientific) was used for data collection. On a BioQuantum K3 direct detection camera (Gatan) using a 20 eV filter slit width operated in zero-loss mode, micrographs were recorded at a nominal magnification of 81,000×, with a calibrated pixel size of 1.095 Å and a defocus range of -0.8 to -2.0 μm. Each movie was exposed for 2.37 s with a dose rate of 25 e^-^/pixel/second, fractionated into 38 frames, resulting a total dose of 50 e^-^/Å^2^.

### Cryo-EM data processing

Movies were aligned using a MotionCor2-like algorithm (Zheng *et al*, 2017) implemented in RELION v3.1 (Zivanov *et al*, 2020), Contrast transfer function (CTF)-estimation and non-templated particle picking were carried out by Warp v1.09 (Tegunov & Cramer, 2019). 4604239 particles were picked in the S/x1 dataset, 967643 particles were picked in the S/x2 dataset, 82693 particles were picked in the S/x3 dataset, and 1375445 particles were picked in the S/native low-pH dataset. Extracted particles were imported back into RELION. An EM map of SARS-CoV-2 S-R/x2 spike (EMD-11329) (Xiong *et al*., 2020) in closed conformation was filtered to 50 Å resolution as the initial model in the first 3D classification of each dataset. Initial 3D classification using the closed spike as initial model was accomplished at bin2 to remove contaminating particles.

Particles within 3D classes displaying clear secondary structures were pooled and subjected to cleaning by one round of 2D classification. For the S/native low-pH dataset, micrographs were imported into cryoSPARC (Punjani *et al*, 2017) to perform blob picking to obtain a template, then 1375445 particles were picked by template picking. One round of 2D classification were followed by the template picking to remove contaminating particles. Selected particles were imported back into RELION to perform an initial 3D classification using a 60 Å filtered closed conformation SARS-CoV-2 S-R/x2 spike map (EMD-11329) as the initial model. Subsequently, a second round of 3D classification using 50 Å filtered closed conformation SARS-CoV-2 S-R/x2 spike map (EMD-11329) was carried out to classify different conformations in each dataset. Auto-refinement, CTF refinement and Bayesian polishing were performed iteratively on classified subsets of different conformations.

To classify dynamic features in Domain D region of locked and closed spikes, we used a previously described focused classification method (Qu *et al*., 2022) on the S/x1 dataset. Briefly, 576514 and 944224 locked and closed spike particles were symmetry expanded in RELION to obtain 1729542 and 2832672 particles, respectively. A round of in-place 3D classification was carried out using a 30 Å sphere mask around the variable Domain D area using a non-low pass filtered consensus map of the region as the reference. Subsequently, each expanded particle was traced back to its original spike trimer particle. The original trimer particles were classified into locked-1, locked-2, locked-112, locked-122 and an asymmetrical closed conformation with extruding FPPR in one protomer. Those classified classes were subjected to another round of 3D auto-refinement.

Following the final round of 3D auto-refinement, map resolutions were estimated by the 0.143 criterion using the phase-randomization-corrected Fourier shell correlation (FSC) curve calculated between two independently refined half-maps multiplied by a soft-edged solvent mask. Final reconstructions were sharpened and locally filtered in RELION. The data processing procedures were summarized in **Fig. S4**. The estimated B-factors of maps are listed in **Table S1**.

### Cryo-EM structure model building

SARS-CoV-1 spike ectodomain structure (PDB: 5X58) was used as the starting model for building SARS-CoV-1 spike structures of closed conformations. SARS-CoV-2 spike ectodomain structures (PDB: 7XTZ/7XU2) were used as the starting model for building SARS-CoV-1 structures of locked conformations. PDBs were fitted into maps in UCSF Chimera v1.16 (Pettersen *et al*, 2004). Manual model rebuilding was performed in Coot 0.9.8 (Casanal *et al*, 2020). Stereochemistry of the manually built models were optimized using Namdinator (https://namdinator.au.dk) (Kidmose *et al*, 2019). Final models were refined in PHENIX v1.19.1 (Afonine *et al*, 2018) by real space refinement with secondary structure restraints and geometry restraints. All structural figures were generated using UCSF Chimera v1.16.

